# Vegetation increases CH_4_ emissions and methanotroph diversity in marine sediments

**DOI:** 10.64898/2026.02.17.706329

**Authors:** Marion Dolivet-Maréchal, Carlos Palacin-Lizarbe, Henri M.P. Siljanen, Dhiraj Paul, Abigaïl Delort, Jonathan Gervaix, Charline Creuzé des Châtelliers, Sabine Schmidt, Mathis Cognat, David Sebag, Olivier Taugourdeau, Clara Schübert, Nathalie Labourdette, Isabelle Bertrand, Lorenzo Rossi, Xavier Le Roux, Agnès Richaume, Alessandro Florio

## Abstract

Seagrass meadows are key blue carbon (C) ecosystems, storing large amounts of organic C over centuries. Their climate benefits may be reduced by methane (CH₄) emissions, whose microbial and environmental descriptors in *Zostera noltei* meadows, dominant seagrass in North-Western Europe, remain poorly understood. We studied CH₄ fluxes, CH₄-producing and consuming microbial communities and sediment physico-chemical parameters in *Z. noltei* meadows and adjacent bare sediments across seven sites in Arcachon Bay, France. *In situ* CH₄ fluxes were measured at low tide and microbial communities were characterised using targeted metagenomics of three functional genes (*mcrA, mmoX, pmoA*) and quantitative PCR. CH₄ fluxes were higher in vegetated than bare sediments (24.4 ± 2.6 vs. 9.4 ± 0.7 µmol m⁻² d⁻¹). Mixed linear models and random forest analyses identified C accumulation rate and CO₂ flux as the strongest positive descriptors of CH₄ fluxes. Vegetated sediments hosted more diverse methanotrophs, while methanogens showed no habitat differences. Four genera (*mcrA–Methanolobus, mmoX–Methylocella, pmoA–Methylococcus, Methyloglobulus*) emerged as abundant, seagrass-associated, correlated with CH₄ fluxes, and highlighted by models. Functional diversity, especially *pmoA* richness, was a stronger microbial descriptor of CH₄ fluxes than gene abundance or a specific genus. Findings indicate *Z. noltei* meadows enhance C burial and CH₄ emission, with methanotroph diversity potentially mitigating CH₄ emissions. Our results provide the first integrated assessment of CH₄ fluxes and their descriptors in *Z. noltei* meadows, highlighting the intertwined nature of C burial and CH₄ emissions and the need to account for both in blue C climate assessments.

## 1 Introduction

Vegetated coastal ecosystems, such as seagrass meadows, play a critical role in global climate regulation by sequestering large amounts of organic carbon (C_org_) in their sediments over the long-term. Despite occupying only ∼0.1% of the ocean surface, seagrass meadows account for up to 18% of global ocean carbon (C) burial (Kennedy et al., 2010; McLeod et al., 2011). Thus, they are defined as key “blue carbon” habitats due to their high primary productivity and capacity to trap organic-rich particles (Duarte et al., 2013; Fourqurean et al., 2012). Yet, the climate mitigation potential of seagrass meadows has recently been questioned due to their ability to emit methane (CH₄) (Bahlmann et al., 2015; Garcias-Bonet & Duarte, 2017), a potent greenhouse gas (GHG) with a global warming potential 27 times higher (IPCC, 2023) than CO₂. These emissions can offset up to 6.1 ± 0.8% of C burial (Dolivet-Maréchal et al., 2025). However, global CH₄ flux estimates from seagrasses remain highly uncertain due to the lack of knowledge on the abiotic and biotic determinants of these emissions (Tan et al., 2025; Yau et al., 2023).

Methane emissions from seagrass sediments mainly arise from the balance between microbial production and consumption (Yau et al., 2023), which determines whether sediments are sources or sink of this GHG. Methanogens are mostly obligate anaerobic archaea and the primary CH₄ producers in sediments (Conrad, 2020; Schorn et al., 2022). These microorganisms use four main pathways, i.e. hydrogenotrophic, acetoclastic, methylotrophic (Conrad, 2020; Täumer et al., 2022) and methoxydotrophic methanogenesis, the latter being only recently described (Täumer et al., 2022). All pathways converge on the final step catalyzed by methyl-coenzyme M reductase (MCR), encoded by the highly conserved *mcrA* gene, which remains a widely used biomarker to detect and quantify methanogens in environmental samples (Evans et al., 2019; Thauer, 2019). Conversely, CH₄ consumption in seagrass sediments is primarily driven by aerobic methanotrophs, which oxidise CH₄ and thus substantially contribute to CH₄ sink (Laanbroek, 2010; Liang et al., 2025). CH₄ oxidation is catalysed by methane monooxygenases (MMOs), enzymes existing in two forms: particulate (pMMO) and soluble (sMMO). The *pmoA* gene encodes a subunit of pMMO, while *mmoX* encodes a component of sMMO; both genes serve as reliable molecular markers for aerobic methanotrophic communities (Liang et al., 2025). The microbial mechanisms underpinning CH₄ production and consumption in seagrass sediments are still poorly understood (Liang et al., 2025; Schorn et al., 2022). In sediments, recent studies indicate that methanogenic lineages are diverse, including *Methanosaetaceae*, *Methanosarcinaceae*, and *Methanobacteriales* (Chen et al., 2022; Schorn et al., 2022; Täumer et al., 2022) while aerobic methanotrophs are dominated by *Gamma-* and *Alphaproteobacteria* (Liang et al., 2025; Täumer et al., 2022). However, most research has focused on a single functional gene (*mcrA*) or 16S-based community analyses. Within the *Zostera* genus, a few studies on *Z. marina* and *Z. japonica* have quantified *mcrA* and used 16S rRNA sequencing to infer methanotrophic diversity, without directly targeting *pmoA* or *mmoX* (Liu et al., 2021; Tan et al., 2025). Thus, the full functional diversity of CH₄-cycling microbes in *Zostera* sediments remains unknown. To date, no study has simultaneously examined the abundance and diversity indices of the three key CH₄-cycling genes, *mcrA*, *mmoX* and *pmoA* in *Z. noltei* sediments.

CH₄ fluxes in seagrass ecosystems can vary widely across spatial and temporal scales (Maher et al., 2015; Rosentreter et al., 2021), yet there is no overarching general descriptor. Most studies tend to focus on either environmental descriptors, classically organic matter (OM) content and quality, which are often reported as key factors influencing CH₄ fluxes (Al-Haj & Fulweiler, 2020), or microbial community dynamics (Schorn et al., 2022). Few integrate both approaches (Tan et al., 2025). Additionally, comparative microbial analyses of vegetated and adjacent bare sediments remain scarce (Tan et al., 2025), limiting our understanding of seagrass-specific microbial controls on CH₄ fluxes. Furthermore, research has predominantly targeted large tropical seagrass species, whereas temperate dwarf species such as *Zostera noltei*, dominant in North-Western Europe, remain poorly characterised in terms of both CH₄ fluxes and microbial CH₄-cycling communities (Al-Haj & Fulweiler, 2020; Eyre et al., 2023).

In a recent study, on the Arcachon Bay, which hosts the largest *Z. noltei* population in Europe, we showed that CH₄ fluxes were significantly higher in vegetated than in bare sediments (24.4 ± 2.6 vs. 9.4 ± 0.7 µmol m⁻² d⁻¹), with total emissions representing 6.1 ± 0.8% of local C burial (Dolivet-Maréchal et al., 2025). Here, we aim to investigate why CH₄ fluxes differ between vegetated and unvegetated sediments, how they are regulated by environmental descriptors, and how the presence of seagrass structures methanogenic and methanotrophic communities. We hypothesised that CH₄ fluxes would be explained by the quantity and quality of OM stored in sediments, and that higher fluxes in seagrass meadows would be associated with greater diversity and abundance of methanogen-related gene. Using an innovative targeted metagenomic approach and quantitative PCR, we screened for CH₄-related genes (*mcrA*, *mmoX*, *pmoA*) across 83 sediment samples, collected from both seagrass meadows and adjacent bare sediments. This scale of sampling is, to our knowledge, unprecedented for microbial studies in seagrass ecosystems. We combined these gene-based microbial analyses with *in situ* CH₄ flux measurements conducted during low tide, when gas exchange between the sediments and the atmosphere is maximised (Deborde et al., 2010), along with physico-chemical, hydrological and sedimentary parameters. Finally, we integrated all data using two types of modelling approaches, i.e. mixed linear models (MLM) and random forest analysis (RFA), to identify key environmental descriptors of CH₄-cycling. By doing so, this study aims to clarify the contribution of *Z. noltei* meadows to refining C budgets in temperate seagrass ecosystems.

## 2 Materials and Methods

### 2.1 Study site and habitat characteristics

Arcachon Bay is a temperate coastal lagoon located on the southern Atlantic coast of France (44°40′ N, 1°10′ W), covering an area of approximately 174 km² (Figure 1). Subject to mesotidal to macrotidal regimes, the bay is connected to the open ocean via two main inlets and is characterised by semi-diurnal tides that drive strong water exchange. In the Arcachon Bay, *Z. noltei* predominantly forms monospecific meadows in the sheltered intertidal mudflats, and represent the largest *Z. noltei* seagrass beds in Europe (Auby et al., 2011). Their abundance have declined by 44% during the last three decades (Muller et al., 2024) because of extreme weather events including heatwaves and storms (Auby et al., 2011; Gamain et al., 2018). As a result, sediment distribution across the bay is spatially heterogeneous: coarse sands and gravels dominate the main tidal channels, sandy muds accumulate in secondary channels, and finer silts are prevalent in intertidal zones (Cognat et al., 2018).

**Figure 1.**
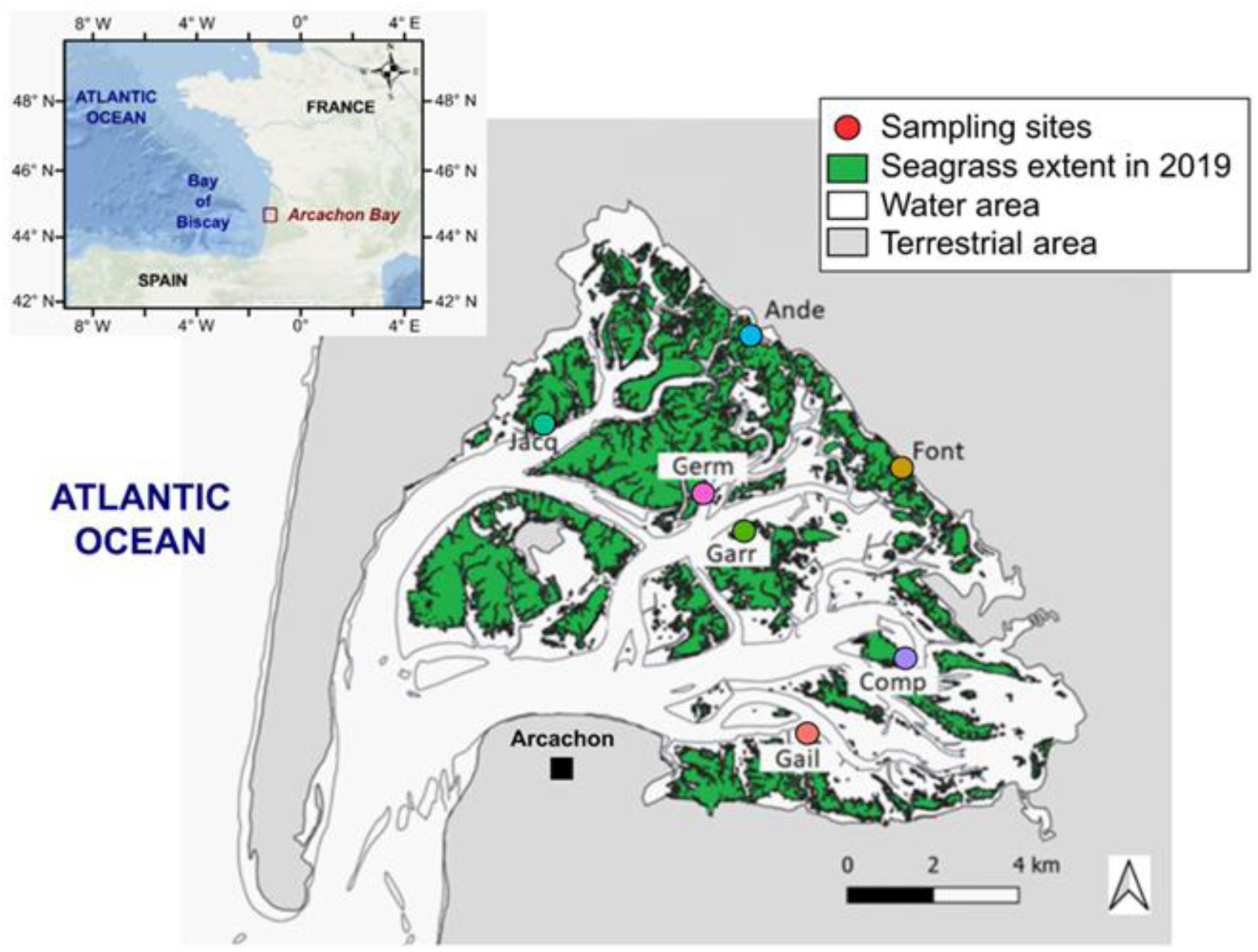
Map of Arcachon Bay with the seven studied sites (colored dots).

### 2.2 Experimental design and sampling strategy

To encompass the environmental diversity of Arcachon Bay, seven sites were selected based on contrasting hydrodynamic regimes, sediment characteristics, and vegetation status (Figure 1). Each site includes two adjacent areas: one vegetated by *Z. noltei* meadows ("Seagrass") and a nearby unvegetated zone ("Bare") (Dolivet-Maréchal et al., 2025), each subsite including six replicates. Historical aerial imagery analyses indicates that these bare areas were vegetated prior to 2005, suggesting potential legacy C storage, with CH₄ production persisting long after plant loss likely sustained by residual methylated compounds from decomposed seagrass material (Schorn et al., 2022). Both types of habitat have remained environmentally stable for over 20 years, providing a consistent framework for comparison.

Two field campaigns were conducted during the weeks commencing September 12, 2022, and August 28, 2023, i.e. during the peak biomass of *Z. noltei* (Auby et al., 2011) (see Dolivet-Maréchal et al., 2025 for details on the sampling design). In 2022, preliminary sediment cores were collected to characterise C burial and screen potential GHG emissions. In 2023, GHG fluxes (CH₄ and CO_2_) were measured *in situ* using opaque static flux chambers, deployed at midday and at low tide, following a standardised layout (5-meter radius). Immediately after each flux measurement, three sediment cores (10 cm depth) were taken at the chamber location and pooled into a single composite sample per replicate, representing the surface sediment layer commonly used for GHG and microbial analyses. A total of 83 flux measurements and composite sediment samples were analysed across the bay (see Dolivet-Maréchal et al., 2025 for site layout and sampling setup). To preserve *in situ* conditions and prevent microbial alteration, samples were stored in dark either at 4°C (for incubations and sediment characterizations) or −20°C (for molecular analyses). Sediment physico-chemical parameters were quantified as described in the Supplementary materials.

### 2.3 *In situ* and laboratory GHG measurements

CH₄ and CO₂ fluxes were measured *in situ* using a LI-COR static Smart Chamber system (model 8200-01S) coupled with a trace gas analyzer (LI-7810; LI-COR Biosciences, USA). The chamber was deployed on pre-installed PVC collars at each replicate point, and measurements were performed over a 3-minute period following a short stabilization phase. Data was recorded every second and processed using LI-COR SoilFluxPro™ software. Flux rates were calculated based on the rate of gas concentration change over time, and expressed in µmol CH₄ m⁻² d⁻¹ or µmol CO₂ m⁻² d⁻¹ (details in Dolivet-Maréchal et al., 2025).

To measure microbial activities, i.e. anaerobic methanogenesis coupled to CO₂ production, aerobic methanotrophy and aerobic CO₂ production, 15 grams of fresh sediment were transferred into airtight flasks sealed with rubber stoppers and incubated at 28°C. Gaz concentrations were measured using a gas chromatograph equipped with a micro-catharometer detector (µGC-R990; SRA instruments, Marcy L’Etoile, France). For methanogenesis and anaerobic CO₂ production, the headspace was flushed with helium to establish anoxic conditions. CH₄ and CO₂ concentrations were measured every 4 hours, starting 6 hours after incubation began, over a 48-hour period. Methanotrophy was assessed using a newly developed protocol under aerobic conditions and with the headspace supplemented with 10% CH₄. Methane concentration was measured every 6 hours, starting 2 hours after incubation, continuing for 74 hours. Aerobic CO₂ production was measured every hour over 6 hours. Microbial activity rates were estimated from the slope of the linear regression of CH₄ (μg C-CH₄ g⁻¹ dry soil h⁻¹) or CO₂ concentration (μg C-CO₂ g⁻¹ dry soil h⁻¹) over time and converted to µmol CH₄ or CO₂ m⁻² d⁻¹.

### 2.4 Abundance of bacteria 16S and functional CH_4_-cycling genes

The abundances of bacterial *16S rRNA* and CH₄-cycling genes (*mcrA*, *mmoX*, *pmoAIa*, *pmoAIb*, and *pmoAII*) were estimated *via* quantitative real-time PCR after DNA extraction and quantification from sediment samples (see full methodological details in Supplementary Materials and Methods and Table S1).

### 2.5 Targeted metagenomic and bioinformatic pipeline

To investigate the functional diversity of CH₄-producing and consuming microorganisms, a targeted metagenomic approach was applied. This method relied on sequence capture by probe hybridization to selectively enrich functional genes involved in CH₄ and mineral nitrogen transformations, including *mcrA*, *mmoX*, and *pmoA*, as described in Putkinen et al., 2021 and Siljanen et al., 2025. Gene-specific probes were designed based on clustered reference sequences at defined similarity thresholds (Siljanen et al., 2025). DNA sequencing was performed using the Illumina platform (Arbor Biosciences, Ann Arbor, MI, USA), producing approximately five million reads per sample. Raw sequences were processed using in-house bioinformatic pipelines on the CSC Puhti supercomputer (Espoo, Finland). Gene sequences were identified using (Eddy, 2022) HMMER v3.3.2 via the nhmmer tool with a threshold of E < 0.0001. Multiple sequence alignments were conducted with MAFFT (Katoh & Standley, 2013), and phylogenetic placements were inferred with RAxML (version 8.2.12) (Stamatakis, 2014). Community composition at the genus level was illustrated using ggplot2 in R (v4.4.0), after merging gene variants assigned to the same genus based on phylogenetic placement. Relative abundances were calculated as the proportion of gene sequences affiliated with each genus within a sample. These values were then multiplied by gene-specific copy numbers per m², as determined by qPCR, to estimate absolute abundances per m². Genera detected for both *mmoX* and *pmoA* genes were labeled with ‘_m’ for *mmoX* and ‘_p’ for *pmoA’* to distinguish between the two.

Alpha diversity metrics were calculated separately for each sample to assess the diversity of *mcrA*, *mmoX*, and *pmoA* gene sequences. Richness (number of unique placements) and Shannon diversity index were computed using the nplace count per phylogenetic placement as a proxy for abundance. These calculations were performed using the R package vegan (v1.44.0) (Oksanen et al., 2020), applying the specnumber() and diversity() functions. In this manuscript, gene-specific richness and diversity are referred to as gene_R and gene_D, respectively (e.g. mcrA_R, pmoA_D).

All sequences are available in the NCBI Sequence Read Archive (PRJNA1261814).

### 2.6 Mixed linear models & random forest analysis

To assess the environmental and microbial descriptors of CH₄ fluxes, MLM were employed. All explanatory variables were log₁₀- or square root-transformed to reduce the impact of extreme values and then scaled (z-scores) to allow direct comparison of their relative influence. Model selection was performed using the dredge function from the MuMIn R package, retaining only models with all significant descriptors (p < 0.05) to avoid overfitting. Model performance was assessed using Corrected Akaike Information Criterion (AICc) and R² values. Linear models were fitted using the lm function, and mixed-effects models with the lme function (R v4.3.1). For fixed-effect models, generalised least squares (gls) models were also fitted to enable direct AICc comparison with mixed-effects models. We tested models with only fixed effects, as well as mixed-effects models including random effects accounting for site variability, sediment type (seagrass vs. bare), or their combination (subsite). For each modelling approach, biotic and abiotic variables were split into different subsets (Table S2A-L).

Separate models were built for each subset using their respective variables to identify key descriptors within each category (Table S2A-L, as in Palacin-Lizarbe et al., 2020). The most significant descriptors selected from each subset-specific model were then merged to create a combined subset (Table S2M). Models based on this combined subset were then constructed to evaluate the overall explanatory power. The most significant models for each subset were selected and are described in the Results section.

In addition, we applied RFA as an alternative modelling approach, using the same subsets of descriptors. This combined modelling method enabled us to identify the most important descriptors driving CH₄ fluxes and to validate the robustness of our findings. It is important to note that RFA does not include the site, type or subsite effects as MLM do. Model tuning was performed using the *tuneRF* function in R, testing different *mtry* values (i.e. the number of variables randomly sampled at each split) across 1000 trees. At each iteration, *mtry* was adjusted by a factor of 1.5, and the search continued only if the relative improvement in out-of-bag (OOB) error exceeded 1%. This procedure enabled the identification of the optimal *mtry* value while minimizing overfitting. By evaluating the mean increase in mean squared error (%IncMSE), key variables within each subset were highlighted, reinforcing the significance of descriptors previously identified in the MLM.

### 2.7 Statistical analyses

All data analyses were conducted using R (version 4.4.0, R Core Team 2024). Results are reported as means ± standard error (SE). Variables were log₁₀- or square root-transformed when necessary to meet analysis assumptions and reduce the influence of extreme values. Differences between bare and vegetated sediments were tested by site and across the entire bay using Wilcoxon tests. To compare gene abundance, richness, and diversity across sites, we performed Kruskal-Wallis tests followed by non-parametric post hoc pairwise comparisons using Dunn’s test, with Benjamini-Hochberg correction for multiple testing. Spearman’s rank correlation test was used to examine relationships between CH₄ fluxes, production and oxidation rates. Only genera significantly correlated with CH₄ fluxes (p < 0.05) and of high abundance were selected for further modelling analyses.

Principal Component Analysis (PCA) was performed to explore patterns among physical, chemical, microbial, and GHG variables across all sampling sites, using the FactoMineR package in R. Additionally, a PCA based on Hellinger distance (Legendre & Gallagher, 2001) was carried out to assess the ordination of microbial gene variants involved in CH_4_-cycling. This analysis aimed to identify genera with gene variants that significantly influence the PCs. Significant correlations between variables and components were identified using the envfit function from the vegan package and incorporated into PCA visualizations.

## 3 Results

### 3.1 Spatial gradients driving sediment functioning

Two main gradients sorted the studied sediments. The first corresponded to a large-scale spatial gradient, while the second was associated with the presence or absence of seagrass at a smaller spatial scale. The large-scale gradient was linked to the connection to the ocean and to the sediment stability, as it was evidenced in the Principal Component 1 (PC1, Figures 2 and S1). On the left side of the PC1 are sorted sites that are more exposed to the hydrodynamic forcing, located near major tidal channels, and closer to the ocean inlet, characterised by longer immersion times, stronger current flows, and greater depths (higher bathymetry). These exposed sites are in more dynamic and physically constrained environments, where organic and mineral C arrive after being carried there by hydrodynamic forces. Conversely, on the right side of PC1 are the sites in the inner part of the bay. These sheltered sites have a lower degree of connection to the ocean and are characterised by calmer conditions, higher levels of labile C, and increased microbial abundance (Figure S1, Table S3).

**Figure 2.**
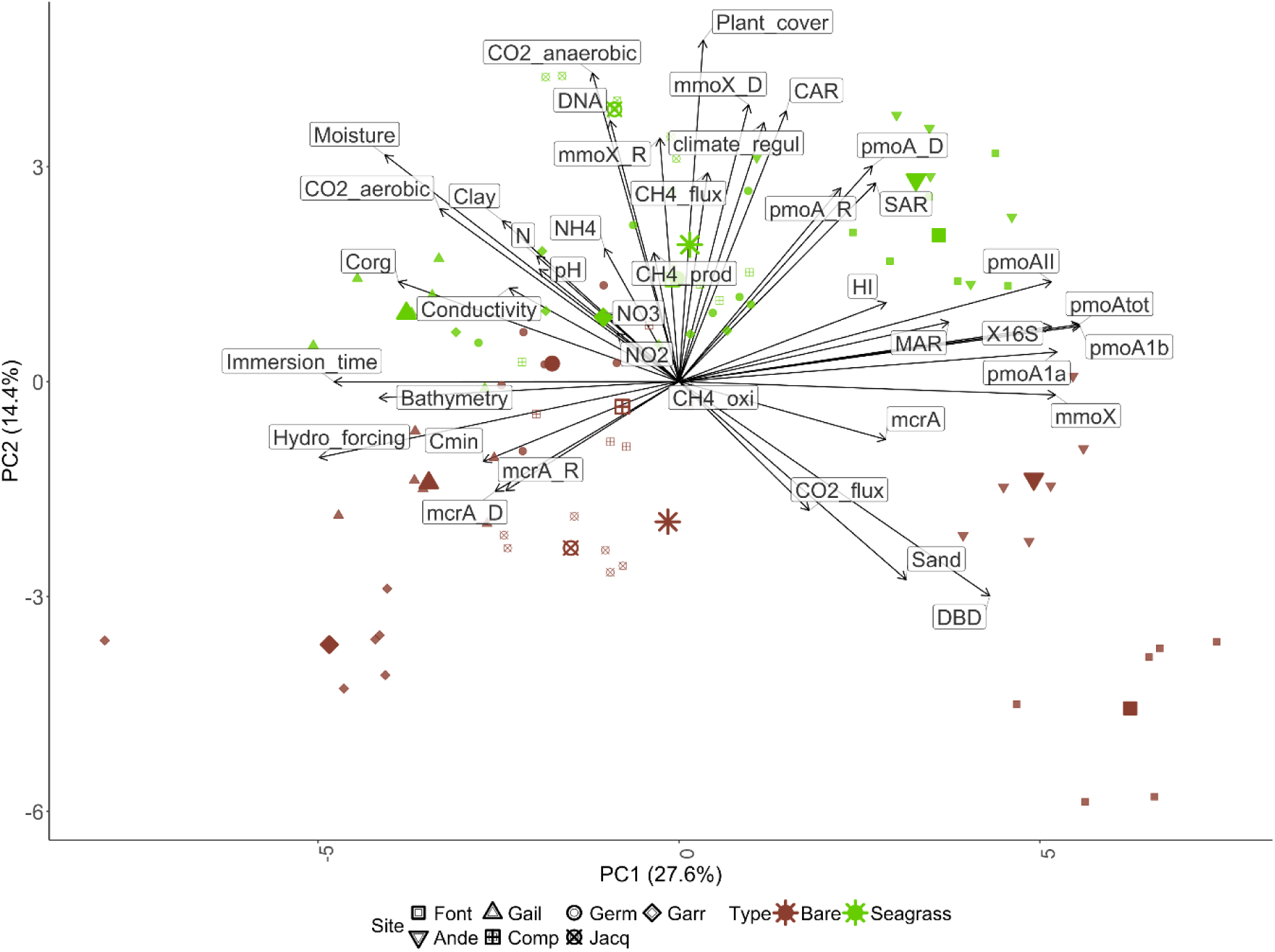
PCA biplot illustrating the relationships among physical, chemical, microbial, and GHG variables across all sampling sites. Arrows represent the contribution of each variable to the ordination. Both axes were rescaled (×6) to improve visualization of sites and variables. Sites are represented by different shapes. Color symbols denote the sediment type (brown = Bare, green = Seagrass), n = 83. Larger symbols represent the barycenter of the subsite, i.e. the combination of site and type.

The second main ecological gradient is driven by the presence of vegetation within sites. Bare sediments are in the negative side of PC2 while vegetated in the positive side (Figure 2). Most of the measured variables tend to have higher values in seagrass-covered areas compared to bare sediments, with the exception of hydrodynamic forcing (Tables 1 and S4). Vegetated subsites are characterised by higher clay content, greater sediment moisture, and elevated values of several biogeochemical variables, including: CH₄ flux, aerobic and anaerobic CO₂ production, CH₄ production, SAR (Sediment Accumulation Rate), MAR (Mass Accumulation Rate), CAR, climate regulation, nitrogen (N), ammonium (NH₄⁺), and nitrate (NO₃⁻) content (Tables 1 and S4). On average across the bay, CO₂ fluxes were negative, indicating net uptake, with no consistent effect of vegetation. However, considerable variability was observed among sites, with two out of seven showing net positive fluxes (Table 1A).

**Table 1.** GHG-related variables (A) and microbial data (B) for the studied sites, grouped by sediment type: bare or vegetated (seagrass). Sites are ordered by increasing CH₄ fluxes measured in seagrass sediments. Significant differences (Wilcoxon tests) between bare and vegetated sediments are indicated as follows: # (p < 0.1), * (p < 0.05), ** (p < 0.01)), *** (p < 0.001), and NS (not significant), n = 83.

| Site_ID | Type<br>(Subsite) | CH<sub>4</sub> flux<br>μmol CH<sub>4</sub> m<sup>-2</sup> d<sup>-1</sup> | CO<sub>2</sub> flux<br>μmol CO<sub>2</sub> m<sup>-2</sup> d<sup>-1</sup> | CO<sub>2</sub> aerobic<br>μmol CO<sub>2</sub> m<sup>-2</sup> d<sup>-1</sup> | CO<sub>2</sub> anaerobic<br>μmol CO<sub>2</sub> m<sup>-2</sup> d<sup>-1</sup> | CH<sub>4</sub> production<br>μmol CH<sub>4</sub> m<sup>-2</sup> d<sup>-1</sup> | CH<sub>4</sub> oxidation<br>μmol CH<sub>4</sub> m<sup>-2</sup> d<sup>-1</sup> |
| --- | --- | --- | --- | --- | --- | --- | --- |
| Comp | Bare | 10.8<sup>NS</sup><br>± 2.2 | -5.5e+04<br>± 8257.3 | 4.0e+05<br>± 5.9e+04 | 1.7e+05<br>± 1.8e+04 | 0.51<br>± 0.06 | 0<sup>NS</sup> |
|  | Seagrass | 13.9<sup>NS</sup><br>± 1.5 | -2.4e+04<sup>**</sup><br>± 4501.9 | 6.6e+05<sup>#</sup><br>± 7.9e+04 | 3.9e+05<sup>**</sup><br>± 6.5e+04 | 4.73<sup>#</sup><br>± 1.73 | 64.03<sup>NS</sup><br>± 64.03 |
| Jacq | Bare | 13.3<sup>NS</sup><br>± 1.1 | -1.6e+04<sup>**</sup><br>± 1825.7 | 5.1e+05<br>± 7.3e+04 | 1.5e+05<br>± 9425.9 | 0.93<sup>NS</sup><br>± 0.31 | 0<sup>NA</sup> |
|  | Seagrass | 14.5<sup>NS</sup><br>± 1.1 | -5.3e+04<br>± 6175.4 | 8.7e+05<sup>**</sup><br>± 5.7e+04 | 2.9e+05<sup>**</sup><br>± 1.7e+04 | 0.57<sup>NS</sup><br>± 0.36 | 0<sup>NA</sup> |
| Germ | Bare | 9.0<br>± 2.0 | -2.4e+04<sup>NS</sup><br>± 7128.5 | 3.7e+05<br>± 1.9e+04 | 1.3e+05<br>± 1.1e+04 | 1.76<sup>NS</sup><br>± 0.67 | 0<sup>NS</sup> |
|  | Seagrass | 15.0<sup>#</sup><br>± 2.2 | -2.2e+04<sup>NS</sup><br>± 7816.4 | 6.4e+05<sup>**</sup><br>± 7.6e+04 | 2.3e+05<sup>**</sup><br>± 1.7e+04 | 0.30<sup>NS</sup><br>± 0.08 | 17.10<sup>NS</sup><br>± 17.10 |
| Font | Bare | 8.1<br>± 0.8 | 1.5e+04<sup>NS</sup><br>± 976.1 | 9.9e+04<br>± 1.4e+04 | 8.1e+04<br>± 7935.9 | 0.03<br>± 0.01 | 0<sup>NS</sup> |
|  | Seagrass | 27.0<sup>**</sup><br>± 4.7 | 1.2e+04<sup>NS</sup><br>± 3184.2 | 5.6e+05<sup>**</sup><br>± 5.1e+04 | 2.9e+05<sup>**</sup><br>± 4.4e+04 | 3.49<sup>**</sup><br>± 1.36 | 66.37<sup>NS</sup><br>± 49.39 |
| Garr | Bare | 11.4<br>± 2.2 | 946.2<sup>NS</sup><br>± 2638.9 | 8.8e+05<sup>#</sup><br>± 4.0e+04 | 1.6e+05<br>± 1.0e+04 | 0.95<br>± 0.18 | 81.71<sup>NS</sup><br>± 81.71 |
|  | Seagrass | 29.7<sup>*</sup><br>± 9.8 | 6736.5<sup>NS</sup><br>± 1987.9 | 7.2e+05<br>± 8.1e+04 | 2.8e+05<sup>**</sup><br>± 2.2e+04 | 2.34<sup>**</sup><br>± 0.41 | 0<sup>NS</sup> |
| Gail | Bare | 4.2<br>± 0.2 | -3.2e+04<sup>NS</sup><br>± 5784.4 | 5.3e+05<sup>NS</sup><br>± 3.5e+04 | 1.4e+05<sup>NS</sup><br>± 1.9e+04 | 0.63<sup>NS</sup><br>± 0.29 | 0<sup>NA</sup> |
|  | Seagrass | 35.0<sup>**</sup><br>± 10.0 | -2.5e+04<sup>NS</sup><br>± 3315.5 | 5.2e+05<sup>NS</sup><br>± 8.3e+04 | 1.9e+05<sup>NS</sup><br>± 2.7e+04 | 2.47<sup>NS</sup><br>± 1.42 | 0<sup>NA</sup> |
| Ande | Bare | 9.0<br>± 2.3 | -2.2e+04<br>± 3612.8 | 2.6e+05<br>± 2.5e+04 | 1.1e+05<br>± 7809.4 | 0.61<sup>NS</sup><br>± 0.42 | 55.15<sup>NS</sup><br>± 48.67 |
|  | Seagrass | 35.9<sup>**</sup><br>± 6.2 | -6106.8<sup>*</sup><br>± 3226.4 | 4.7e+05<sup>*</sup><br>± 7.6e+04 | 2.0e+05<sup>**</sup><br>± 2.0e+04 | 1.53<sup>NS</sup><br>± 0.55 | 31.51<sup>NS</sup><br>± 31.51 |
| All | Bare | 9.4<br>± 0.7 | -1.8e+04<sup>NS</sup><br>± 3629.0 | 4.4e+05<br>± 3.9e+04 | 1.3e+05<br>± 6159.7 | 0.78<br>± 0.15 | 20.03<sup>NS</sup><br>± 13.82 |
|  | Seagrass | 24.4<sup>***</sup><br>± 2.6 | -1.6e+04<sup>NS</sup><br>± 3618.4 | 6.4e+05<sup>***</sup><br>± 3.2e+04 | 2.7e+05<sup>***</sup><br>± 1.6e+04 | 2.20<sup>*</sup><br>± 0.43 | 25.57<sup>NS</sup><br>± 12.43 |

B
| Site_ID | Type<br>(Subsite) | DNA<br>ng m<sup>-2</sup> | 16S<br>copies m<sup>-2</sup> | mcrA<br>copies m<sup>-2</sup> | mmoX<br>copies m<sup>-2</sup> | pmoA1a<br>copies m<sup>-2</sup> | pmoA1b<br>copies m<sup>-2</sup> | pmoAll<br>copies m<sup>-2</sup> | pmoAtot<br>copies m<sup>-2</sup> | mcrA_R | mcrA_D | mmoX_R | mmoX_D | pmoA_R | pmoA_D |
| --- | --- | --- | --- | --- | --- | --- | --- | --- | --- | --- | --- | --- | --- | --- | --- |
| Comp | Bare | 5.1e+08<br>± 9.9e+07 | 1.4e+14<br>± 1.1e+13 | 6.3e+09<br>± 1.2e+09 | 1.3e+10<sup>NS</sup><br>± 1.6e+09 | 7.7e+10<sup>NS</sup><br>± 1.1e+10 | 7.6e+11<sup>NS</sup><br>± 9.7e+10 | 6.3e+10<sup>NS</sup><br>± 1.2e+10 | 9.0e+11<sup>NS</sup><br>± 1.2e+11 | 25.8<sup>NS</sup><br>± 4.3 | 2.8<sup>NS</sup><br>± 0.1 | 37.2<sup>NS</sup><br>± 1.8 | 3.1<sup>NS</sup><br>± 0.1 | 10.4<sup>NS</sup><br>± 1.5 | 0.4<br>± 0.0 |
|  | Seagrass | 1.2e+09**<br>± 1.8e+08 | 2.0e+14#<br>± 1.9e+13 | 2.1e+10#<br>± 6.8e+09 | 1.1e+10<sup>NS</sup><br>± 1.7e+09 | 6.5e+10<sup>NS</sup><br>± 9.2e+09 | 9.2e+11<sup>NS</sup><br>± 1.2e+11 | 7.6e+10<sup>NS</sup><br>± 1.4e+10 | 1.1e+12<sup>NS</sup><br>± 1.4e+11 | 27.3<sup>NS</sup><br>± 1.8 | 2.8<sup>NS</sup><br>± 0.1 | 33.8<sup>NS</sup><br>± 1.7 | 3.1<sup>NS</sup><br>± 0.0 | 8.8<sup>NS</sup><br>± 0.9 | 0.6#<br>± 0.1 |
| Jacq | Bare | 3.8e+08<br>± 8.7e+07 | 9e+13<sup>NS</sup><br>± 9.4e+12 | 1.5e+10#<br>± 3.4e+09 | 6.8e+09<sup>NS</sup><br>± 3.6e+08 | 3.8e+10<sup>NS</sup><br>± 3.5e+09 | 5.1e+11<br>± 3.9e+10 | 3.4e+10<sup>NS</sup><br>± 1.9e+09 | 5.8e+11<br>± 4.3e+10 | 57.5<sup>NS</sup><br>± 18.5 | 3.1<sup>NS</sup><br>± 0.3 | 23.7<br>± 1.4 | 2.4<br>± 0.0 | 7.3<sup>NS</sup><br>± 1.4 | 0.3<sup>NS</sup><br>± 0.1 |
|  | Seagrass | 2.9e+09**<br>± 6.9e+08 | 1e+14<sup>NS</sup><br>± 1.5e+13 | 7.0e+09<br>± 1.6e+09 | 7.6e+09<sup>NS</sup><br>± 1.3e+09 | 4.3e+10<sup>NS</sup><br>± 5.2e+09 | 6.7e+11#<br>± 6.0e+10 | 4.7e+10<sup>NS</sup><br>± 6.8e+09 | 7.6e+11*<br>± 7.1e+10 | 20.7<sup>NS</sup><br>± 2.4 | 2.7<sup>NS</sup><br>± 0.1 | 32.3*<br>± 1.3 | 3.0**<br>± 0.0 | 6.2<sup>NS</sup><br>± 0.9 | 0.5<sup>NS</sup><br>± 0.1 |
| Germ | Bare | 1.4e+09<sup>NS</sup><br>± 2.5e+08 | 7.9e+13<br>± 1.0e+13 | 9.3e+09<br>± 1.0e+09 | 9.8e+09<br>± 1.5e+09 | 5e+10<br>± 4.9e+09 | 6.6e+11<br>± 6.4e+10 | 4.6e+10<br>± 6.0e+09 | 7.5e+11<br>± 7.3e+10 | 27.5<sup>NS</sup><br>± 3.0 | 2.9*<br>± 0.1 | 28.8<sup>NS</sup><br>± 2.8 | 2.9<sup>NS</sup><br>± 0.1 | 7.5<sup>NS</sup><br>± 0.7 | 0.5<sup>NS</sup><br>± 0.1 |
|  | Seagrass | 1.6e+09<sup>NS</sup><br>± 2.4e+08 | 1.9e+14*<br>± 3.5e+13 | 2.3e+10*<br>± 4.8e+09 | 1.7e+10*<br>± 2.8e+09 | 1e+11#<br>± 1.9e+10 | 1.2e+12#<br>± 2.1e+11 | 9.7e+10*<br>± 1.5e+10 | 1.4e+12#<br>± 2.4e+11 | 22.3<sup>NS</sup><br>± 2.5 | 2.6<br>± 0.0 | 30.7<sup>NS</sup><br>± 2.0 | 3.0<sup>NS</sup><br>± 0.1 | 7.0<sup>NS</sup><br>± 0.7 | 0.6<sup>NS</sup><br>± 0.1 |
| Font | Bare | 5.6e+08<br>± 5.7e+07 | 3.4e+14<sup>NS</sup><br>± 6.1e+13 | 2.3e+10<br>± 9.6e+09 | 3.2e+10**<br>± 3.5e+09 | 1.5e+11#<br>± 2.0e+10 | 2.0e+12<sup>NS</sup><br>± 3.1e+11 | 1.3e+11<sup>NS</sup><br>± 2.8e+10 | 2.3e+12<sup>NS</sup><br>± 3.5e+11 | 21.2<br>± 2.6 | 2.7<sup>NS</sup><br>± 0.1 | 20.3<br>± 2.6 | 2.6<br>± 0.1 | 4.7<br>± 1.3 | 0.5<br>± 0.2 |
|  | Seagrass | 7.6e+08#<br>± 7.7e+07 | 2.7e+14<sup>NS</sup><br>± 3.6e+13 | 4.5e+10#<br>± 4.9e+09 | 1.5e+10<br>± 2.8e+09 | 9.7e+10<br>± 1.9e+10 | 1.5e+12<sup>NS</sup><br>± 1.3e+11 | 9.8e+10<sup>NS</sup><br>± 1.4e+10 | 1.7e+12<sup>NS</sup><br>± 1.6e+11 | 34.3#<br>± 4.3 | 2.7<sup>NS</sup><br>± 0.1 | 29.3*<br>± 1.9 | 3.0*<br>± 0.1 | 15.8**<br>± 2.2 | 1.1#<br>± 0.2 |
| Garr | Bare | 5.1e+08<br>± 7.6e+07 | 3.4e+13<br>± 4.9e+12 | 1.7e+10<sup>NS</sup><br>± 2.0e+09 | 3.9e+09<sup>NS</sup><br>± 5.5e+08 | 2.0e+10<br>± 3.0e+09 | 2.8e+11<br>± 4.4e+10 | 1.4e+10<br>± 2.1e+09 | 3.1e+11<br>± 4.8e+10 | 59.7#<br>± 3.8 | 3.7*<br>± 0.1 | 26.3<br>± 1.8 | 2.6<br>± 0.1 | 6.2<br>± 1.2 | 0.3<br>± 0.1 |
|  | Seagrass | 1.2e+09**<br>± 3.7e+08 | 1.0e+14**<br>± 1.7e+13 | 2.2e+10<sup>NS</sup><br>± 8.5e+09 | 7.2e+09<sup>NS</sup><br>± 1.7e+09 | 4.2e+10#<br>± 7.4e+09 | 6.4e+11*<br>± 1.1e+11 | 5.3e+10#<br>± 1.6e+10 | 7.3e+11*<br>± 1.3e+11 | 47.3<br>± 5.2 | 3.4<br>± 0.0 | 30.8#<br>± 2.7 | 2.8#<br>± 0.1 | 10.0*<br>± 0.9 | 0.6*<br>± 0.1 |
| Gail | Bare | 1.8e+09<sup>NS</sup><br>± 4.1e+08 | 6.6e+13<sup>NS</sup><br>± 4.5e+12 | 1.8e+10*<br>± 6.9e+09 | 6.0e+09<sup>NS</sup><br>± 8.1e+08 | 3.1e+10<sup>NS</sup><br>± 3.7e+09 | 4.6e+11<sup>NS</sup><br>± 4.6e+10 | 2.4e+10<sup>NS</sup><br>± 5.1e+09 | 5.1e+11<sup>NS</sup><br>± 5.3e+10 | 35.8<br>± 3.5 | 2.7<br>± 0.1 | 26.5<sup>NS</sup><br>± 1.1 | 2.9<sup>NS</sup><br>± 0.1 | 4.2<sup>NS</sup><br>± 0.7 | 0.5<sup>NS</sup><br>± 0.1 |
|  | Seagrass | 1.3e+09<sup>NS</sup><br>± 3.0e+08 | 5.8e+13<sup>NS</sup><br>± 7.8e+12 | 8.5e+09<br>± 8.8e+08 | 4.8e+09<sup>NS</sup><br>± 4.7e+08 | 2.7e+10<sup>NS</sup><br>± 2.8e+09 | 3.6e+11<sup>NS</sup><br>± 3.5e+10 | 2.2e+10<sup>NS</sup><br>± 2.8e+09 | 4.1e+11<sup>NS</sup><br>± 4.0e+10 | 64.7*<br>± 14.3 | 3.4**<br>± 0.2 | 27.7<sup>NS</sup><br>± 2.9 | 2.9<sup>NS</sup><br>± 0.1 | 6.7<sup>NS</sup><br>± 1.4 | 0.6<sup>NS</sup><br>± 0.1 |
| Ande | Bare | 8.0e+08<br>± 7.2e+07 | 2.1e+14<sup>NS</sup><br>± 2.5e+13 | 5.9e+10**<br>± 8.9e+09 | 2.5e+10#<br>± 2.9e+09 | 9.5e+10<sup>NS</sup><br>± 8.5e+09 | 1.6e+12*<br>± 1.1e+11 | 1e+11<sup>NS</sup><br>± 1.3e+10 | 1.8e+12#<br>± 1.3e+11 | 38.7**<br>± 2.9 | 2.9<sup>NS</sup><br>± 0.2 | 26.8<br>± 0.8 | 2.9<br>± 0.0 | 12.0<sup>NS</sup><br>± 2.1 | 0.7<br>± 0.1 |
|  | Seagrass | 1.2e+09#<br>± 1.8e+08 | 1.7e+14<sup>NS</sup><br>± 3.5e+13 | 1.6e+10<br>± 3.0e+09 | 1.6e+10<br>± 2.8e+09 | 7.6e+10<sup>NS</sup><br>± 8.7e+09 | 1.2e+12<br>± 1.2e+11 | 1e+11<sup>NS</sup><br>± 1.4e+10 | 1.4e+12<br>± 1.4e+11 | 23.3<br>± 1.9 | 2.7<sup>NS</sup><br>± 0.0 | 34.0*<br>± 2.0 | 3.1#<br>± 0.1 | 17.0<sup>NS</sup><br>± 1.9 | 1.5**<br>± 0.1 |
| All | Bare | 8.5e+08<br>± 1.0e+08 | 1.4e+14<sup>NS</sup><br>± 1.8e+13 | 2.2e+10<sup>NS</sup><br>± 3.3e+09 | 1.4e+10<sup>NS</sup><br>± 1.7e+09 | 6.6e+10<sup>NS</sup><br>± 7.5e+09 | 9.0e+11<sup>NS</sup><br>± 1.1e+11 | 5.9e+10<br>± 7.7e+09 | 1.0e+12<sup>NS</sup><br>± 1.2e+11 | 38.3<sup>NS</sup><br>± 3.5 | 3.0<sup>NS</sup><br>± 0.1 | 26.9<br>± 1.0 | 2.8<br>± 0.0 | 7.4<br>± 0.6 | 0.5<br>± 0.0 |
|  | Seagrass | 1.5e+09***<br>± 1.6e+08 | 1.6e+14<sup>NS</sup><br>± 1.4e+13 | 2.1e+10<sup>NS</sup><br>± 2.5e+09 | 1.1e+10<sup>NS</sup><br>± 1.0e+09 | 6.5e+10<sup>NS</sup><br>± 5.9e+09 | 9.3e+11<sup>NS</sup><br>± 7.3e+10 | 7.1e+10#<br>± 6.3e+09 | 1.1e+12<sup>NS</sup><br>± 8.4e+10 | 34.3<sup>NS</sup><br>± 3.2 | 2.9<sup>NS</sup><br>± 0.1 | 31.2***<br>± 0.8 | 3.0***<br>± 0.0 | 10.2**<br>± 0.8 | 0.8***<br>± 0.1 |

### 3.2 Key parameters on CH_4_-cycling microbial communities

The abundance and diversity of the CH_4_-cycling genes showed different patterns depending on sediment type and site location. Sediments located in sheltered sites far from the ocean inlet hosted the more abundant CH_4_-cycling prokaryotic communities with higher abundances of *pmoA*, *mmoX*, and *mcrA* (also for the overall bacterial communities), while vegetated sediments hosted more diverse methanotrophic communities (Figure 2, Tables 1B and S3A). In all studied sediments, *pmoA* was the more abundant functional gene (1.0^12^ ± 7.3^10^ copies per m²) followed by *mcrA* (2.1^10^ ± 2.1^09^ copies per m²) and by *mmoX* (1.2^10^ ± 1.0^09^ copies per m²), (p < 0.001). The diversity and richness followed a different order being higher in *mcrA* followed by *mmoX* and by *pmoA* (p < 2 × 10⁻¹⁶, Tables 1B and S3B, Figure S2).

In vegetated sediments, we observed higher CH₄ fluxes, higher total DNA content, and distinct CH₄-cycling microbial communities (Table 1A-B). For *mcrA*, 7 of 36 genera showed significant associations, 5 enriched in seagrass (i.e. *Methanohalophilus, Mathanolubus, Methanolobus, Methanobacterium, Methanocaldococcus*) and 2 enriched in bare sediments (i.e. *Methanocalculus, Methanoregula*) (Figure 3A). For *mmoX*, 6 of 18 genera were enriched in seagrass (i.e. *Methylibium, Mycobacterium, Methylocaldum, Methylomagnum, Methylococcus, Methylocella*), while none were specific to bare sediments (Figure 3B). These genera were significantly enriched in vegetated compared to bare sediments, although they did not represent the most abundant taxa within the community. In contrast, among the 17 *pmoA* genera, 5 (i.e. *Methylococcus, Methylobacter, Methyloglobulus, Methylomicrobium, Methylomonas*) were both abundant and significantly associated with seagrass (Figure 3C).

**Figure 3.**
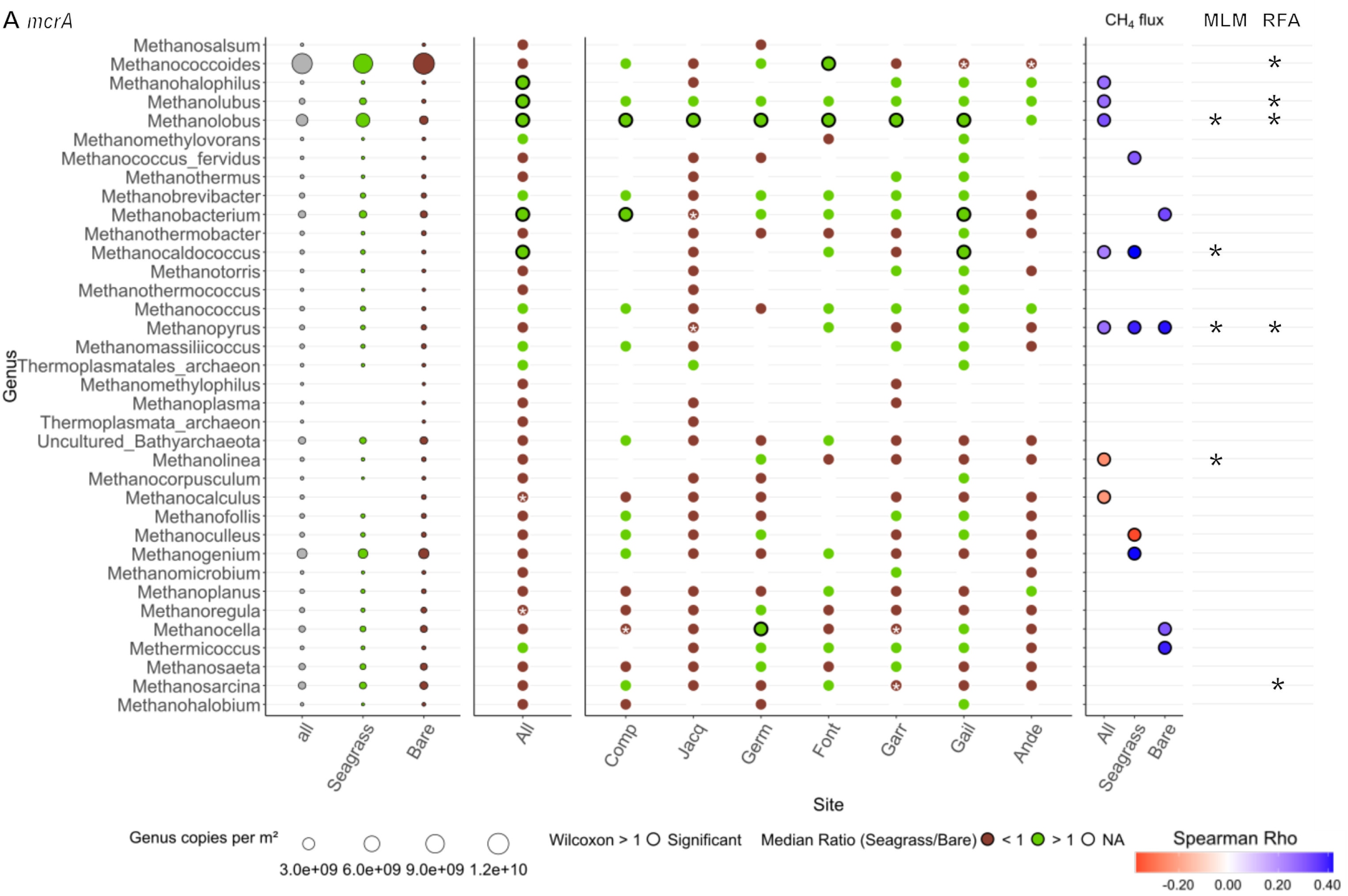
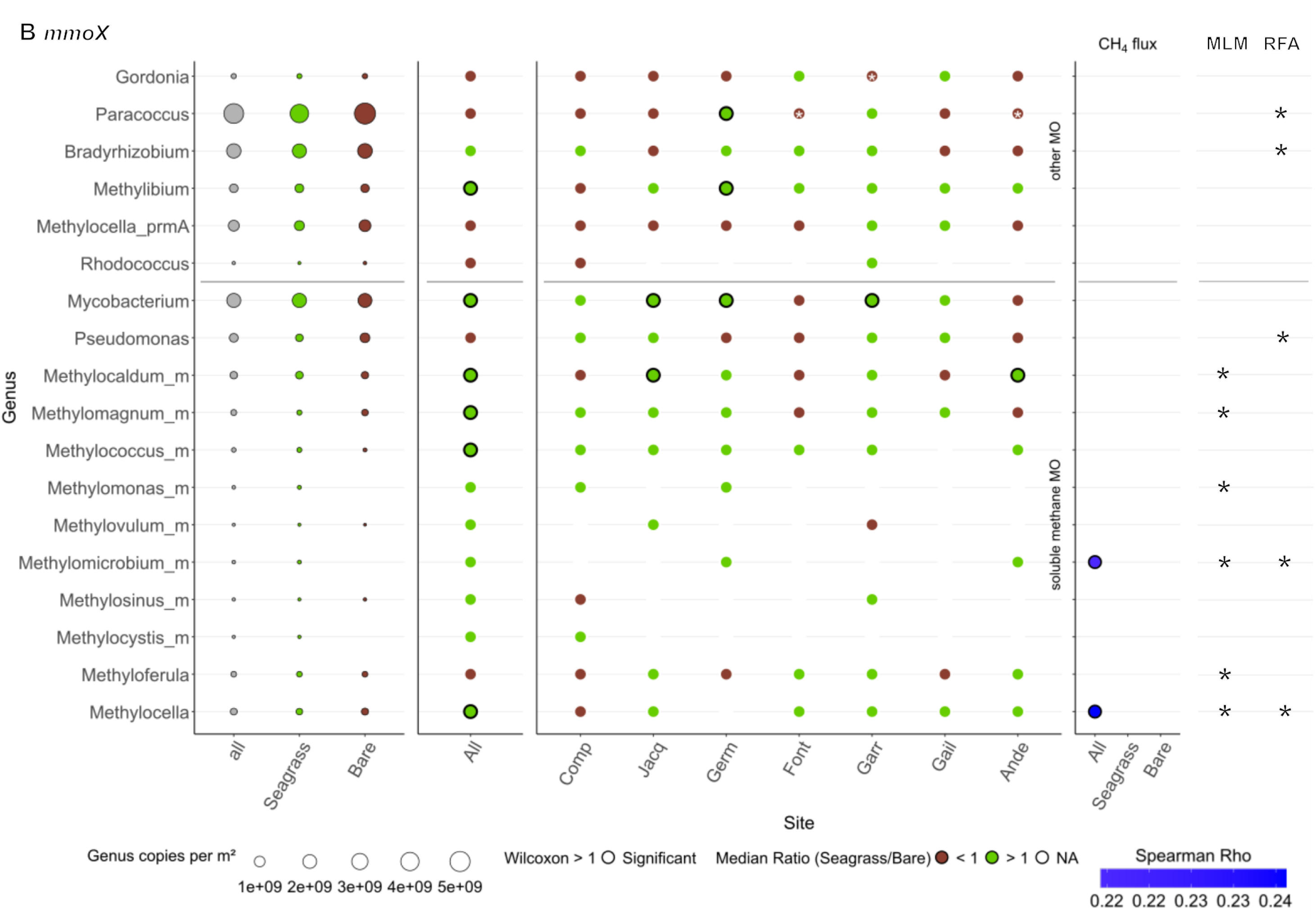
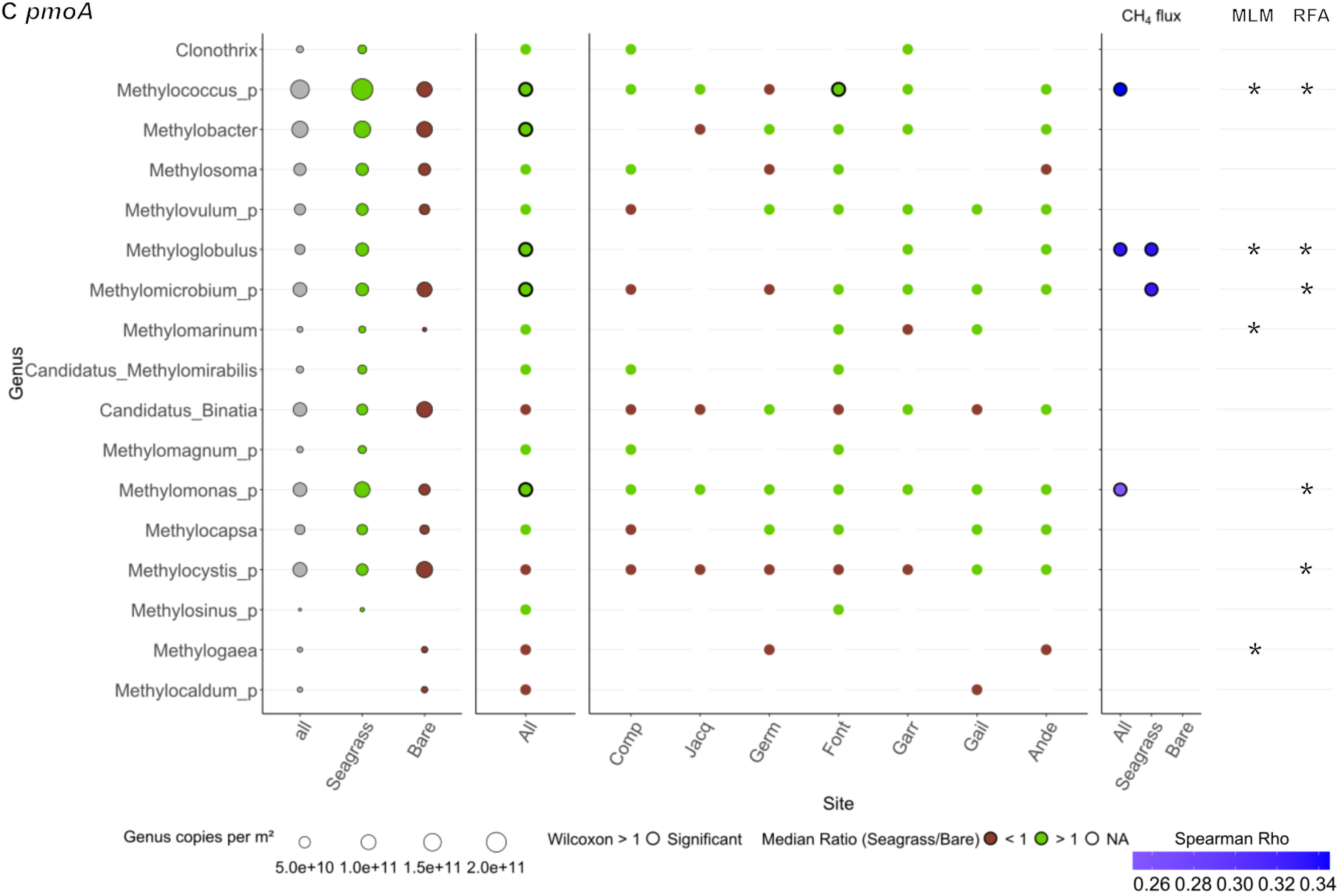
Differential abundance and relationships with CH₄ flux for methanogenic and methanotrophic genera across seagrass and bare sediments in Arcachon Bay. Panel A includes genera identified from the *mcrA* gene (methanogenesis marker), panel B from *mmoX*, and panel C from *pmoA* (methanotrophy markers). For each genus, the first three columns show mean abundance (gene copies per m², scaled by point size) across all samples, seagrass, and bare sediments, n = 83. The next two panels display, respectively, overall and site-specific median abundance ratios (Seagrass/Bare), color-coded (green: ratio > 1, seagrass affinity; brown: ratio < 1, bare affinity). Sites are ordered by increasing CH₄ fluxes measured in seagrass sediments. Significant differences (Wilcoxon test, *P* < 0.05) are indicated by a black circle for seagrass preference and a white star within the brown point for bare preference. The penultimate panel shows only significant Spearman correlations (Rho) between genus abundance and CH₄ flux, with colors indicating correlation strength and direction (blue: positive, red: negative). The final panel displays genera identified as important markers of CH₄ flux based on mixed linear models (MLM) and random forest analysis (RFA), indicated by asterisks. “NA” denotes genera absent in both habitats.

Ordination of microbial communities based on absolute genus abundances (Figure 4) mirrored the global PCA combining environmental, microbial, and GHG variables (Figure 2), with vegetation presence and related environmental variables emerging as the main structuring factors. Only a subset of genera correlated significantly with CH₄ fluxes (Figure 3), and even fewer were specifically linked to CH₄ production or oxidation (Figure S3). Moreover, the genera correlated with each of these processes were generally not the same. Four genera (*mcrA–Methanolobus, mmoX–Methylocella, pmoA–Methylococcus, Methyloglobulus*) consistently emerged as key taxa (Figures 3 and 4), being abundant, seagrass-associated, correlated with CH₄ fluxes, and supported by MLM and RFA (marked with asterisks in Figure 3). However, the whole community (or its diversity) rather than specific genera seems to be marking the CH4 flux (details in the next section).

**Figure 4.**
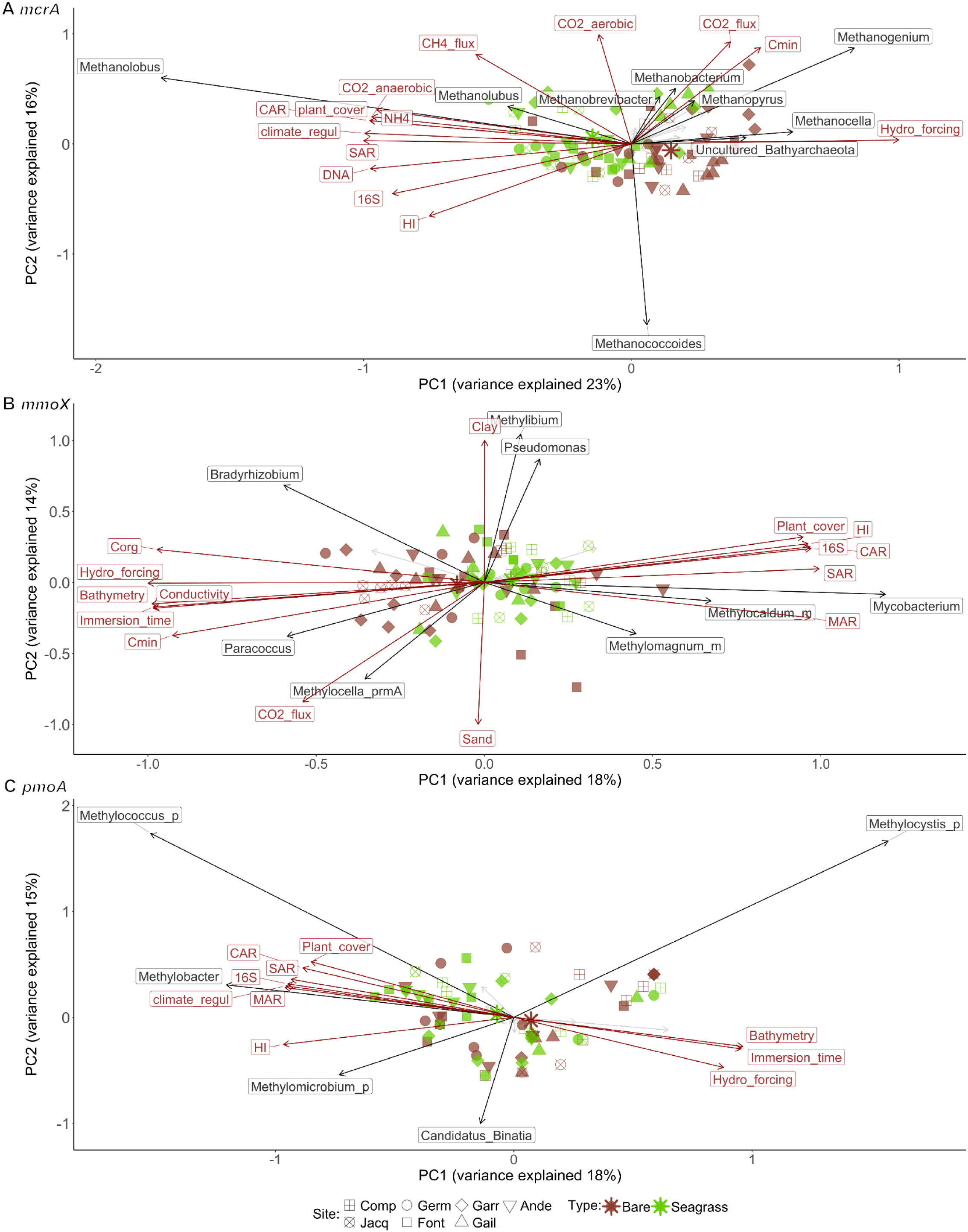
Principal Component Analysis (PCA) biplots illustrating the relationships between microbial genera (based on functional genes) and environmental variables across all sampling sites. The analyses are based on the absolute abundance of genera associated with the genes: (A) *mcrA*, (B) *mmoX*, and (C) *pmoA*. Only microbial genera with significant contributions to PC1 and PC2 are displayed in black, and only environmental variables with significant correlation with components are shown in red. Sites are represented by different shapes, and symbol colors denote sediment type, n = 83. Larger star symbols indicate the barycenter of each type.

### 3.3 Modelling of CH₄ Fluxes

Models of CH_4_ fluxes in response to spatial variability, gas, physical, chemical and microbial sediment descriptors, as well as all descriptors combined, explain up to 88% of the variation observed (Table 2). About the spatial variability, only the presence of seagrass explains 60% of the CH₄ fluxes (model 12). Adding the site variability in combination with the seagrass presence, i.e. subsite variability (model 7), just increases the explanatory power to 64%, while the site variability alone does not describe fluxes at all (0%, model 22).

**Table 2.** Summary of the best significant mixed linear models explaining methane (CH₄) fluxes across different environmental conditions. Models are ranked by Corrected Akaike Information Criterion (AICc), from lowest (best fit) to highest, and are grouped according to fixed or random factors: site, sediment type (bare vs. seagrass), and subsite. Within each group, the top-performing models are shown in bold. All models include CH₄ flux as the response variable, with physical, gas-related, chemical, or microbial variables used as predictors. Only the best model(s) per subset are presented here and n = 83.

| Model | Formula: Fixed part | Random part | AICc (gls) | Fixed R<sup>2</sup> | Global R<sup>2</sup> | Subset |
| --- | --- | --- | --- | --- | --- | --- |
| 1 | $\log_{10}(\text{CH}_4\_flux) = 0.95 \times \text{CAR} - 0.56 \times \text{MAR} + 0.43 \times \text{CO}_2\_flux$ | ~1\|Subsite | 186 | 0.43 | 0.77 | Combined |
| 2 | $\log_{10}(\text{CH}_4\_flux) = 0.64 \times \text{CAR} - 0.39 \times \text{MAR} + 0.33 \times \text{CO}_2\_flux - 0.29 \times \log_{10}(\text{DNA})$ | ~1\|Type | 186 | 0.21 | 0.65 | Combined |
| 3 | $\log_{10}(\text{CH}_4\_flux) = 0.35 \times \text{CO}_2\_flux$ | ~1\|Subsite | 190 | 0.11 | 0.69 | Gas |
| 4 | $\log_{10}(\text{CH}_4\_flux) = 0.48 \times \text{CAR}$ | ~1\|Subsite | 192 | 0.22 | 0.65 | Chemical |
| 5 | $\log_{10}(\text{CH}_4\_flux) = 1.66 \times \text{CAR} - 1.28 \times \text{Hydro\_forcing} - 1.27 \times \text{SAR} + 0.77 \times \text{Bathymetry} + 0.46 \times \text{CO}_2\_flux$ | ~1\|Site | 193 | 0.57 | 0.88 | Combined |
| 6 | $\log_{10}(\text{CH}_4\_flux) = 0.25 \times \text{sqrt}(\text{pmoA\_R}) - 0.24 \times \text{mmoX\_D} - 0.22 \times \log_{10}(\text{DNA})$ | ~1\|Type | 193 | 0.09 | 0.76 | Microbial |
| 7 |  | ~1\|Subsite | 194 | 0 | 0.64 | Subsite |
| 8 | $\log_{10}(\text{CH}_4\_flux) = 0.25 \times \text{CO}_2\_flux$ | ~1\|Type | 196 | 0.05 | 0.64 | Gas |
| 9 | $\log_{10}(\text{CH}_4\_flux) = 0.27 \times \text{CAR} - 0.22 \times \text{HI}$ | ~1\|Type | 199 | 0.07 | 0.59 | Chemical |
| 10 | $\log_{10}(\text{CH}_4\_flux) = 1.30 \times \text{CAR} - 0.72 \times \text{SAR}$ | ~1+CAR\|Site | 200 | 0.28 | 0.80 | Chemical |
| 11 | $\log_{10}(\text{CH}_4\_flux) = 0.21 \times \text{Sand}$ | ~1\|Type | 200 | 0.03 | 0.67 | Physical |
| 12 |  | ~1\|Type | 201 | 0 | 0.60 | Type |
| 13 | $\log_{10}(\text{CH}_4\_flux) = 1.10 \times \text{CAR} - 0.72 \times \text{MAR} + 0.51 \times \text{CO}_2\_flux + 0.27 \times \log_{10}(16S) - 0.20 \times \text{HI}$ | | 183 (201) | 0.53 | | Combined |
| 14 | $\log_{10}(\text{CH}_4\_flux) = -1.74 \times \text{Hydro\_forcing} + 0.94 \times \text{Bathymetry} - 0.78 \times \text{DBD}$ | ~1+DBD\|Site | 202 | 0.50 | 0.88 | Physical |
| 15 | $\log_{10}(\text{CH}_4\_flux) = 0.72 \times \text{CAR} - 0.32 \times \text{MAR}$ | | 216 (224) | 0.26 | | Chemical |
| 16 | $\log_{10}(\text{CH}_4\_flux) = 0.54 \times \text{CAR} - 0.22 \times \text{HI}$ | | 216 (225) | 0.26 | | Chemical |
| 17 | $\log_{10}(\text{CH}_4\_flux) = 0.44 \times \log_{10}(\text{CO}_2\_anaerobic) + 0.34 \times \text{CO}_2\_flux$ | | 217 (225) | 0.25 | | Gas |
| 18 | $\log_{10}(\text{CH}_4\_flux) = 0.44 \times \log_{10}(\text{CO}_2\_anaerobic) + 0.34 \times \text{CO}_2\_flux$ | ~1\|Site | 228 | 0.27 | 0.28 | Gas |
| 19 | $\log_{10}(\text{CH}_4\_flux) = 0.41 \times \text{sqrt}(\text{pmoA\_R})$ | | 226 (231) | 0.16 | | Microbial |
| 20 | $\log_{10}(\text{CH}_4\_flux) = -0.57 \times \text{Hydro\_forcing} - 0.46 \times \text{DBD}$ | | 224 (232) | 0.18 | | Physical |
| 21 | $\log_{10}(\text{CH}_4\_flux) = 0.41 \times \text{sqrt}(\text{pmoA\_R})$ | ~1\|Site | 234 | 0.16 | 0.16 | Microbial |
| 22 |  | ~1\|Site | 243 | 0 | 0.00 | Site |

About gas and chemical descriptors, sediments with high long-term C sequestration (CAR) and high CO_2_ fluxes show also high CH₄ fluxes. These two variables (CAR and CO_2_ flux) are included in the best models, either fixed models (13, explaining up to 53%), or mixed models including the subsite variability (models 1, 3, and 4, explaining up to 77%) and the effect of vegetation (model 2 explaining up to 65%).

About the physical variables, model 14 (explaining up to 88%) includes the site variability and shows negative effects of the hydrodynamic forcing and of sediment dry bulk density, the latter related to the higher density in bare sediments, and positive effect of bathymetry, pointing to higher fluxes in deeper sites. Model 11 includes the positive effect of sand with the presence of seagrass.

The best microbial model includes a positive association with *pmoA* richness, and negative associations with *mmoX* diversity and DNA content (model 6, explaining up to 76%). The negative coefficients do not imply a causal inhibition but reflect the partial effects of these variables combined with *pmoA* richness. Diversity metrics are more informative than functional gene abundance (qPCR) or genus-level composition, with the *pmoA* gene emerging as the most influential descriptor (Figure 5E, Table S2D-L, and Figure S4A-B).

**Figure 5.**
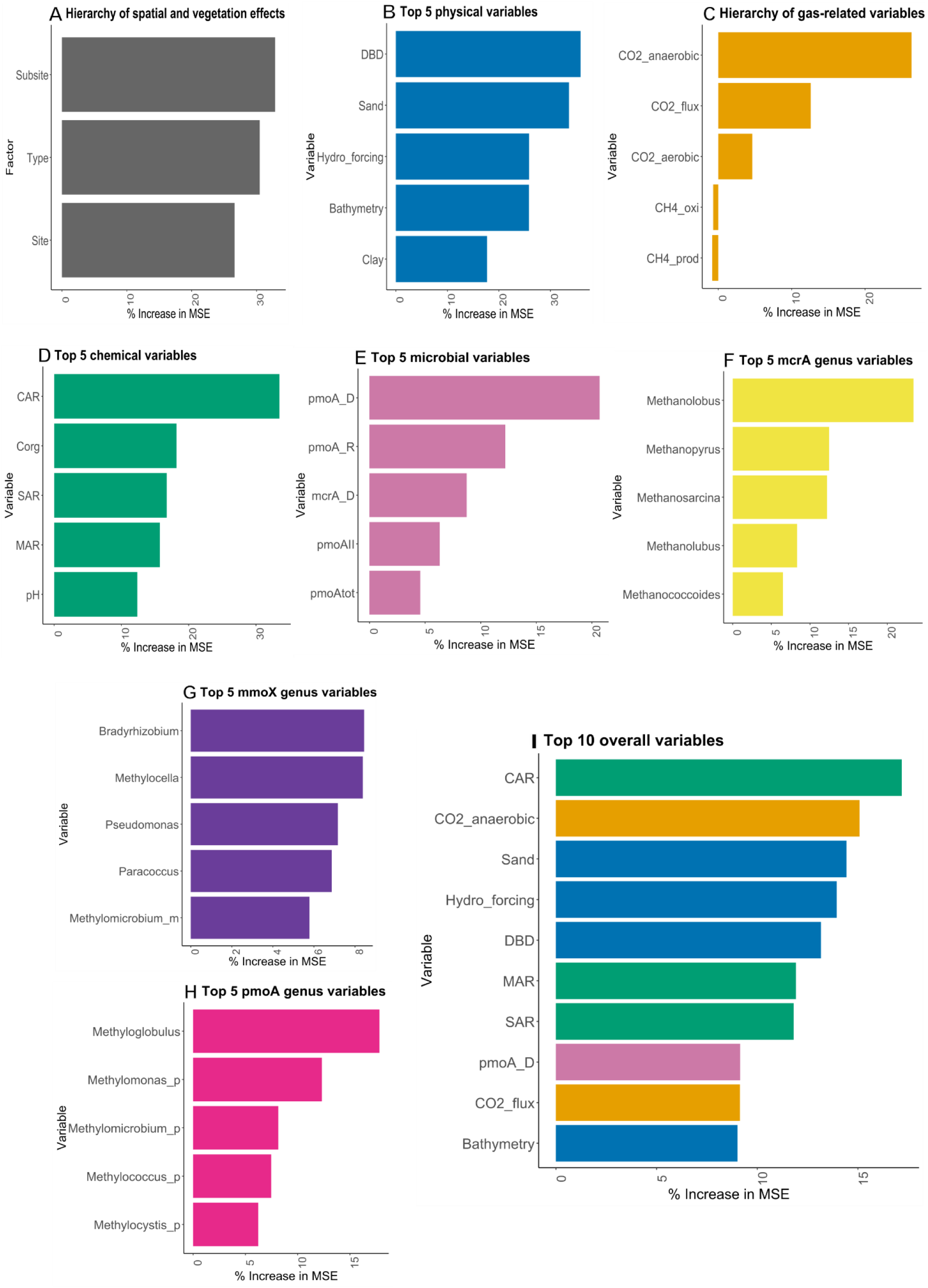
Variable importance in random forest analysis identifying key descriptors of methane (CH₄) fluxes. (A) Hierarchy of spatial and vegetation effects (site, sediment type, and subsite). (B-D) Top 5 predictors within each environmental category: (B) physical, (C) gas-related, and (D) chemical variables, n = 83. (E) Top 5 microbial predictors based on functional gene abundances, diversity indices, *16S* copies, and DNA content. (F-H) Top 5 taxonomic predictors at the genus level for *mcrA*, *mmoX*, and *pmoA* genes, respectively. (I) Overall top 10 predictors of CH₄ flux across all variables.

The best models combine variables from different subsets, i.e. gas, chemical, physical and microbial subsets, always including the positive effect of CAR and CO_2_ fluxes and the negative of MAR or SAR (models 1, 2, 5 and 13). The best model (1) includes the subsite variability, and the effect of CAR, MAR and CO_2_ flux. The best model (2) includes the effect of vegetation and has the same descriptors of model 1 adding the negative effect of the DNA content. Model 5 includes the site variability, and the effect of MAR is not included but SAR, hydrodynamic forcing, and bathymetry are included instead. Finally, the best fixed model (13) is like model 1 adding *16S* copies and C lability (HI). The RFA overall agrees with the models confirming high-influencing variables (Figure 5I) which are included in several models.

## 4 Discussion

### 4.1 Low CH_4_ fluxes in Arcachon bay compared to other seagrass meadows

We observed higher CH_4_ fluxes in seagrass-vegetated areas compared to bare sediments, agreeing with previous studies (Al-Haj & Fulweiler, 2020; Tan et al., 2025). Two recent syntheses comparing CH₄ fluxes across the sediment-water-air continuum in seagrass ecosystems reported median values of 64.8 and 35.8 µmol CH₄ m⁻² d⁻¹, respectively (Al-Haj & Fulweiler, 2020; Eyre et al., 2023). In comparison, we observed an average flux of 24.4 ± 2.6 µmol CH₄ m⁻² d⁻¹ in vegetated zones, with a median of 13.7 µmol CH₄ m⁻² d⁻¹, is lower than both values but still indicates a significant contribution of seagrass sediments to atmospheric CH₄ emissions. The measured CH_4_ fluxes represent 6.1 ± 0.8% of the estimated C burial in the Arcachon Bay (Dolivet-Maréchal et al., 2025). These results highlight the need to consider both long-term sequestration and emission processes when evaluating the climate regulation potential of blue C habitats.

Our measurements were performed at midday during low tide, using opaque chambers on exposed sediments to standardize sampling across sites. While this approach likely captured near-maximum emission rates, it does not fully account for diel or tidal variability, which can influence CH₄ release through changes in sediment oxygenation, water coverage, and temperature (Deborde et al., 2010; Maher et al., 2015; Rosentreter et al., 2021). In some temperate seagrass meadows, CH₄ fluxes peak in the afternoon and evening rather than at night, suggesting that thermal and physical drivers can partially override redox-dependent diel patterns (Henriksson et al., 2024). Thus, while our measurements provide robust estimates of relative differences between vegetated and bare sediments, continuous in situ monitoring would be valuable to capture short-term dynamics and further refine flux estimates.

Seagrass meadows in our study area have declined by approximately 50% since the 1990s (Muller et al., 2024). As restoration efforts to increase seagrass distribution may enhance CH₄ emissions alongside C burial, future studies should explore how meadow age, species, coverage, and biomass per area may influence flux variability. Moreover, a dedicated study on N₂O emissions, using a similar approach to ours, would also be valuable, given its high warming potential (273 times greater than CO_2_) (IPCC, 2023) and the current lack of data (Murray et al., 2015).

### 4.2 C-related features as key descriptors of CH₄ fluxes

The combined use of MLM and RFA identified C-related variables as the most consistent and important descriptors of CH₄ fluxes across our sites, supporting our initial hypothesis that CH₄ fluxes are influenced by sedimentary OM, particularly the C related traits. CAR and CO₂ flux positively influenced CH₄ emissions, while SAR, MAR, and the lability of OM (HI) showed negative associations.

These results suggest that C-rich environments with more recalcitrant OM tend to support higher CH₄ emissions. This is possibly due to more C in deeper sediments, where anaerobic conditions are conducive to methanogenesis and sustained microbial activity. The negative relationship with lability (HI) supports the idea that labile OM is preferentially degraded through aerobic or alternative anaerobic pathways, reducing CH₄ production (Le Mer & Roger, 2001).

Also, the particle size matters, as finer particles and more C_org_, promote both C storage and CH₄ emissions (Al-Haj & Fulweiler, 2020; Eyre et al., 2023). This can be explained, in part, by a positive feedback loop in seagrass ecosystems: the presence of vegetation slows down water flow, enhancing the deposition of fine particles and organic matter. This further stabilises the sediments (Cognat, 2019) and fosters anaerobic conditions favourable to methanogenesis despite the O_2_-extrusion in the roots. Indeed, our best-performing model with site as a random factor, highlighted the negative role of hydrodynamic forcing in modulating CH₄ fluxes, agreeing with Jin et al., 2025. Stronger hydrodynamics increases sediment mixing which disrupts anaerobic microsites and favours CH₄ oxidation or alternative heterotrophic metabolism, and could even increases sediment instability till avoiding seagrass presence and C burial.

Importantly, the presence of seagrass alone explained a large proportion (60%) of the variance in CH₄ fluxes. This suggests that vegetation not only facilitates C accumulation but also drives biogeochemical processes through its influence on sediment characteristics and microbial communities.

Overall, our results indicate that while increased C burial represents a clear benefit for climate regulation, it is accompanied by a disservice in the form of higher CH₄ emissions. Nevertheless, these processes appear synergistic, as sites with greater C burial also tend to exhibit higher CH₄ emissions. Overall, our results suggest that and CH₄ emissions are not in a trade-off relationship. Instead, they appear to co-occur, as areas with higher C burial also tend to exhibit higher CH₄ emissions. All C-related variables, except HI, were positively associated with CH₄ fluxes, indicating a cumulative effect of C quantity on both sequestration and emission. Therefore, promoting C storage in these systems may inherently involve accepting increased CH₄ emissions, a conclusion supported by both our statistical models and the PCA. This dual role of vegetation, enhancing both C sequestration and CH₄ production, is consistent with earlier observations (Dolivet-Maréchal et al., 2025; Eyre et al., 2023; Kristensen et al., 2025; Macreadie et al., 2019) and reinforces the complexity of blue C system dynamics.

### 4.3 Diversity of methanotrophs is the best microbial CH_4_ flux descriptor

Modelling revealed that functional diversity of CH_4_-cycling genes was a better descriptor of CH₄ fluxes than functional gene abundance or than the presence of specific genus of microbes carrying CH_4_-cycling genes. The most explanatory microbial model identified *pmoA* richness as a positive descriptor, while *mmoX* diversity and total DNA content were negatively associated with CH₄ emissions, reflecting their partial effects in the context of the multivariate model. Also, we observed significant and positive correlations between each of the *pmoA* and *mmoX* genera and CH_4_ fluxes, these genera being more abundant in vegetated than bare sediments. These findings suggest that diverse methanotrophic communities in CH₄-rich environments may help buffer CH₄ release by enhancing oxidation capacity (Arnold et al., 2023; Knief, 2015; Tan et al., 2025).

Our results show that microbial communities differ between bare and seagrass sediments, with the key genera highlighted in this study being more abundant in vegetated sites. In addition, methanotroph diversity is higher in vegetated sediments. However, contrary to our initial hypothesis, the abundance and diversity indices of methanogens were not significantly higher in vegetated sediments compared to bare sediments. Furthermore, *mcrA* variables were not included in models of CH_4_ fluxes, agreeing with other studies with no clear links between methanogens and CH_4_ production potential (Berberich et al., 2020; Chaudhary et al., 2017). This may reflect the fact that methanogenesis can significantly occur also at deeper anoxic layers, which were not included in our 0-10 cm sediment sampling. A more stratified and deeper sampling approach will be needed to help distinguish vertical patterns in community structure and activity. In contrast, we observed higher diversity and richness of *pmoA-* and *mmoX-*methanotrophs in seagrass sediments compared to unvegetated ones, a pattern also reported by Tan et al., 2025. Vegetation likely promotes the development of diverse methanotrophic communities by enhancing sediment oxygenation via root O₂ release (Brodersen et al., 2024) and photosynthetic O₂ production, both of which can stimulate CH₄ oxidation (Lyimo et al., 2018).

Although taxonomic identity alone explained less variation in CH₄ fluxes than functional diversity, certain genera stood out. *Methanolobus* (*mcrA*) was particulary notable, being also more abundant in vegetated than in bare sediments in the Chinese coast (Tan et al., 2025) and contributing to CH₄ production in brackish Baltic Sea sediments (Tsola et al., 2024), consistent with our observations showing a positive correlation with *in situ* CH₄ fluxes. Aerobic methanotrophic genera such as *Methylococcus* and *Methylomicrobium*, which harbor both *mmoX* and *pmoA* genes and belong to the *Gammaproteobacteria* class, have also been reported to be strongly associated with seagrass sediments (Tan et al., 2025). *Methylococcus* has been identified as a predominant aerobic methanotroph in marine sediments (Håvelsrud et al., 2011), in line with our results showing it as the most abundant genus for the *pmoA* gene. In addition, *Methylocella*, a facultative methanotroph capable of using CH₄ as well as multicarbon substrates such as acetate, pyruvate, and malate has been observed across a wide range of ecosystems, including lake and marine sediments (Rahman et al., 2011), and was also detected in our marine sediments, showing a preference for vegetated areas and a positive correlation with *in situ* CH₄ fluxes. Similarly, *Methyloglobulus* has been associated with aquatic rhizospheres and identified as one of the most abundant methanotrophs carrying the *pmoA* gene, consistent with our observations in vegetated sediments, suggesting a potential role in CH₄ oxidation within root-associated aquatic zones (Bao et al., 2025). These findings reinforce the idea that seagrass habitats support complex and spatially coupled CH₄ production and oxidation processes.

Our approach enabled a comprehensive characterization of key CH₄-cycling microbial groups, but some technical limitations remain. For qPCR, the *pmoA* gene was targeted with three primer sets, whereas *mmoX* and *mcrA* were each amplified with a single primer set, which may have contributed to the higher *pmoA* abundance observed. Similarly, reference databases are more complete for *mcrA*-associated methanogens than for methanotrophs, potentially affecting richness and diversity estimates, although *mmoX* values were also high despite the smaller database. These limitations highlight the importance of balanced primer design and curated databases (Campbell et al., 2022). Future studies integrating RNA- or protein-based analyses could identify metabolically active microbes and provide functional insights into *in situ* CH₄-cycling (Markovski et al., 2025).

## 5 Conclusion

This study provides new insights into the biogeochemical functioning of seagrass ecosystems by showing that CH₄ emissions, although generally modest, are highly influenced by C levels. Seagrass presence enhances CH₄ fluxes through organic C accumulation, finer sediment particles and reduced hydrodynamic forcing. Vegetated sediments also showed more diverse methanotrophic communities. These CH₄ emissions can co-occur with high blue C sequestration, reflecting a complex ecological trade-off rather than a simple dichotomy. As such, seagrass restoration and management strategies should consider both aspects of their climate impact. To better quantify this balance, future research should integrate high-resolution flux monitoring with activity-based microbial profiling.

## Acknowledgements

We thank Ifremer, PNMBA, and Pierrick Fribourg for providing nautical resources. We are also grateful to the two interns, Rémi Dugué and Margot Ahr, for their valuable assistance during the fieldwork and laboratory analyses.

## Competing interests

The authors declare no competing interests.

## Funding

This work was supported by the Green Deal project REST-COAST, funded by the EU Horizon2020 programme (Grant Agreement 101037097). Additional support was provided by the Graduate School H2O’Lyon (ANR-17-EURE-0018) of Université de Lyon (UdL), within the program *“Investissements d’Avenir”* operated by the French National Research Agency (ANR).

## Data and code availability

The data used to calculate the MAR are archived on the SEANOE platform [https://doi.org/10.17882/99990]. The Excel file and R code, which include the other variables used in this study and those for figure generation, are archived on Zenodo at [https://doi.org/10.5281/zenodo.17243268]. The Excel file contains data collected from seven sites within Arcachon Bay, and the R code includes statistical analyses (e.g. Wilcoxon tests, mixed linear models, and random forest analyses) as well as scripts for figure generation. Additionally, raw targeted metagenomic sequences generated in this study have been deposited in the NCBI Sequence Read Archive (SRA) under BioProject accession number PRJNA1335804.

## Notes

https://doi.org/10.17882/99990

https://doi.org/10.5281/zenodo.17243268

### Competing Interest Statement

The authors have declared no competing interest.

