## Supplementary_methods_figures for "Vegetation increases CH_4_ emissions and methanotroph diversity in marine sediments"

**Supplementary text**

**Materials and Methods**

**Characterisation of the sediments and sites**

**Plant cover**

Seagrass (*Zostera noltei*) coverage was quantified following the European Water Framework Directive protocol (Auby et al., 2018). At each study point, a 30 × 30 cm quadrat was placed, photographed, and the percentage of seabed covered by seagrass was independently estimated by three researchers. The final coverage value for each quadrat was calculated as the average of these three estimates.

**Hydrodynamic forcing**

Hydrodynamic forcing was assessed using a dimensionless parameter, which quantifies the total hydrodynamic energy during a tidal cycle (spring or neap tide) that exceeds a critical velocity threshold for sediment erosion or leaf uprooting. This parameter was calculated following sediment transport model formulations (Le Hir et al., 2011), and applies to both vegetated and unvegetated areas. Further methodological details are available in Dolivet-Maréchal et al., 2025.

**Analyses on the top 10 cm of mixed sediment cores**

**Abundance of bacterial 16S and functional $CH_4$-cycling genes**

DNA was extracted from sediment samples using the NucleoSpin® Soil Kit (Macherey-Nagel GmbH & Co, Düren, Germany) (Béraud et al., 2025) following the manufacturer's protocol. DNA concentrations were quantified using a Qubit® 2.0 fluorometer and the Quant-iT™ dsDNA Broad Range (BR) Assay Kit (Invitrogen, France). The abundance of the bacterial community was estimated by absolute quantification of *16S rRNA* gene copy number via quantitative real-time PCR (qPCR), using iTaq™ Universal SYBR® Green Supermix (Bio-Rad) on a CFX Connect

Real-Time PCR Detection System (Bio-Rad). The abundance of key genes involved in $CH_4$ production, i.e. *mcrA* (methanogenic archaea), and $CH_4$ consumption, i.e. *mmoX* (soluble methane monooxygenase), and three distinct clades of *pmoA* (particulate methane monooxygenase: *pmoAIa*, *pmoAIb*, and *pmoAII*) were also quantified. Detailed information on thermal conditions and primer sets is provided in Table S1. Melting curve analysis was conducted to verify PCR product specificity after amplification. Amplification showed a linear response ($R^2 > 0.99$) with no detectable PCR inhibitors, which were systematically checked.

**Moisture**

Moisture content of sediment samples was determined following ISO 11465 (gravimetric method: drying at 105 °C for 24 h and mass difference). Although this standard originates from soil sciences, it has been applied successfully in marine contexts (Shakeel et al., 2022), ensuring comparability and robustness of measurement.

**pH**

Sediment pH was measured using a protocol adapted from ISO 10390:2005. A 4 g portion of lyophilized sediment was placed into a 50 mL Falcon® tube, and 20 mL of deionized water were added. The suspension was mechanically shaken for 1 hour at 140 rpm and 20 °C. Samples were then left to rest at room temperature for 1 to 3 hours before measurement. After manual homogenization and pH meter calibration, the pH value was recorded either after 2 minutes or once the reading had stabilized.

**Conductivity**

Sediment conductivity was assessed following a procedure inspired by ISO 11265:1994. A 20 g aliquot of lyophilized sediment was placed into a 200 mL plastic flask, to which 100 mL of deionized water were added. The suspension was then vortexed vigorously five times at 30-minute intervals over a 2.5-hour period. After agitation, the mixture was filtered using a

Whatman No. 5 filter, and conductivity was measured on the resulting filtrate using a calibrated conductivity meter.

#### Sediment grain size

The sediment grain size distribution was measured using a Malvern laser diffraction instrument (LS13320XR), covering particle sizes from 0.4 to 2000 µm. Sediment samples (a few milligrams) were pretreated by removing organic matter with hydrogen peroxide and dispersed using sodium hexametaphosphate before analysis. The instrument's high sensitivity allowed small sample volumes to be used, ensuring accurate and efficient granulometry. The distribution of particle sizes was expressed as the percentage of clay (<2 µm), silt (2-50 µm), and sand (50-2000 µm).

#### Elementary analyser, nitrogen content

Total nitrogen (N) content was quantified using elemental analysis (e.g. Carlo Erba NA-1500 CNS analyzer) on decarbonated, dried, unsalted and homogenized sediment samples. The analyzer combusts the material at high temperature (1200°C), allowing precise measurement of nitrogen as a percentage of dry weight.

#### Nitrogen analyses

For organic nitrogen pool analysis, fresh soil samples (5 g dry weight equivalent) were transferred into 150 mL plasma vials. After determining the soil moisture content, a 0.01 M $CaCl_2$ solution (Houba et al., 2000) was added. The vials were sealed with parafilm (M®), and the soil suspensions were incubated at 20 °C with shaking (140 rpm) for 2 hours at 10 °C. Following incubation, the suspensions were filtered using membranes with a 0.22 µm pore size. Concentrations of $NO_3^-$, $NO_2^-$, and $NH_4^+$ were determined using a Smartchem 200 sequential analyzer (AMS Alliance, Villeneuve-la-Garenne, France).

#### Rock-Eval® Thermal analyses, hydrogen index

Rock-Eval® thermal analysis is a geochemical technique widely used to quantify and characterize the carbon content of sediment samples (Baudin et al., 2015). In this method, a small, dried, unsalted, and homogenized sediment sample undergoes two successive ramped heating steps: the first under an inert atmosphere (pyrolysis), followed by a second under an oxidative atmosphere.

During the pyrolysis phase, the sample is gradually heated from 200°C to 650°C. The thermal cracking of organic compounds releases a series of volatile products, mainly hydrocarbon effluents, carbon monoxide (CO), and carbon dioxide ($CO_2$), which are continuously monitored by flame ionization detector (FID) and infrared (IR) sensors. In the subsequent oxidation phase, the residual sample is heated from 200°C to 850°C in the presence of oxygen. This step enables the combustion of the remaining organic matter and the thermal decomposition of the mineral fraction, again releasing CO and $CO_2$, which are measured using IR detectors.

Based on the type, quantity, and temperature of gas release, Rock-Eval® allows the estimation of both total organic carbon (TOC [% g C·$g^{-1}$]) and mineral carbon (MinC [% g C·$g^{-1}$]). However, TOC values may sometimes be biased, particularly in non-carbonaceous samples, due to the misattribution of certain carbon signals, leading to an overestimation of the mineral carbon content. To correct for this analytical artifact, the SOTHIS correction method (Hazera et al., 2023; Sebag et al., 2022) was applied to the entire dataset. In the remainder of this article, *Cmin* refer to the MinC values corrected using the SOTHIS method.

In addition to quantitative carbon measurements, Rock-Eval® also provides qualitative indicators such as the Hydrogen Index (HI [mg HC·$g^{-1}$ TOC]). HI is calculated as the ratio between the quantity of hydrocarbons released during pyrolysis (S2 peak) and the TOC content (HI = S2/TOC × 100, in mg HC·$g^{-1}$ TOC). Higher HI values generally indicate hydrogenrich, well-preserved (typically autochthonous) organic matter, while lower values reflect more degraded or terrestrially derived inputs.

**Analyses on the top 20 cm of undisturbed sediment cores**

Sediment cores (up to 20 cm depth) were manually collected using 35 cm long PVC tubes inserted until resistance. In the lab, the cores were sectioned at 1 cm intervals, dried at 62 °C; wet and dry weights were used to calculate the dry bulk density (DBD), assuming a mineral density of 2.65 g $cm^{-3}$. The Sediment Accumulation Rate (SAR) (cm $yr^{-1}$) and Mass Accumulation Rate (MAR) (g $cm^{-2}$ $yr^{-1}$) were calculated using excess $^{210}Pb$ ($Pb_{xs}$) profiles in the sediment, following Amann et al., 2023. A Broad-Energy (@Mirion) gamma spectrometer was used to measure $^{210}Pb$, $^{226}Ra$, and $^{232}Th$ at the EPOC laboratory. The excess of 210Pb ($210Pb_{xs}$), incorporated in sediments from atmospheric inputs and whose decrease with depth provides access to time, was obtained by subtracting $^{226}Ra$ from total $^{210}Pb$ and normalized using $^{232}Th$. Sediment ages were estimated using the Constant Initial Concentration model (Robbins & Edgington, 1975). The SAR and MAR correspond to the slope of the depth-age and cumulative mass-age relationships, respectively.

$C_{org}$ was measured in composite samples prepared from sediment layers located below the surface horizon affected by early diagenesis. Surface sediments were excluded to avoid bias due to organic matter degradation occurring within the uppermost centimeters. Following Delgard et al., 2013, who showed in Arcachon Bay that beyond 3-5 cm depth %C remains relatively constant down to 20 cm, we considered this deeper section as the “stable interval.” Composite samples were prepared by pooling the selected layers within this interval for each core, with each layer weighted by its dry bulk density (DBD) to account for differences in sediment mass per unit volume. Sediments were ground, sieved (2 mm), and decarbonated following Lorrain et al., 2003. $C_{org}$ content was determined using a Thermo FlashSmart

Elemental Analyzer at the EPOC laboratory. CAR (g C $m^{-2}$ $yr^{-1}$) was calculated as the product of MAR and $\%C_{org}$, and converted to $CO_2$ equivalents using a 44:12 molar ratio. The net climate benefit of each habitat was assessed by balancing CAR against GHG emissions. Further details on this method are available in Dolivet-Maréchal et al., 2025. These values (SAR, MAR and CAR) were calculated for the sediment layer corresponding to the last 17-year, the recent period of sedimentary stabilization.

**Supplementary tables**

**Table S1** Target genes, primer names and sequences, amplicon sizes, and references used for quantitative PCR. Total pmoA (pmoAtot) abundance was calculated by summing the gene copy numbers of the three pmoA clades, providing a comprehensive estimate of particulate methane monooxygenase gene abundance in the samples.

| Target Gene | Primer Name | Sequence (5'→3') | Amplicon Size (bp) | Thermal Conditions | Reference |
|---|---|---|---|---|---|
| ***mcrA*** | mcrA_F_mod_SLB | GGTGTMGGATTCACMCARTAYGCWAC | 500 | Denaturation: 95°C/15min<br>- 39 cycles: 95°C/15s; 57°C/45s; 72°C/45s; 78°C/5s (data acquisition)<br>- Melting curve analysis: 0.5°C/0.05s from 65°C to 95°C | Modified by S. Brauer for detecting *M. boonei*, original Hales ME2; also modified by Luton et al. (2002) |
| | mcrA_R_mod_SLB | TTCATTGCRTAGTTHGGRTAGTT | | | |
| ***mmoX*** | mmoXLF | GAAGATTGGGGCGGCATCTG | 427 | Denaturation: 95°C/3min<br>- 39 cycles: 95°C/15s; 63°C/30s; 80°C/5s (data acquisition)<br>- Melting curve analysis: 0.5°C/0.05s from 65°C to 95°C | Rahman et al. (2011) |
| | mmoXLR | CCCAATCATCGCTGAAGGAGT | | | |
| ***pmoAIa*** | A189f | GGNGACTGGGACTTCTGG | 491 | Denaturation: 95°C/3min<br>- 39 cycles: 95°C/15s; 60°C/30s; 72°C/30s; 80°C/5s (data acquisition)<br>- Melting curve analysis: 0.5°C/0.05s from 65°C to 95°C | Kolb et al. (2003) |
| | mb601R | ACRTAGTGGTAACCTTGYAA | | | |
| ***pmoAIb*** | A189f | GGNGACTGGGACTTCTGG | 491 | Denaturation: 95°C/3min<br>- 39 cycles: 95°C/15s; 60°C/30s; 72°C/30s; 80°C/5s (data acquisition)<br>- Melting curve analysis: 0.5°C/0.05s from 65°C to 95°C | Kolb et al. (2003) |
| | mc468R | GCSGTGAACAGGTAGCTGCC | | | |
| ***pmoAII*** | II223F | CGTCGTATGTGGCCGAC | 491 | Denaturation: 95°C/3min<br>- 39 cycles: 95°C/15s; 67°C/30s; 72°C/30s; 82°C/5s (data acquisition)<br>- Melting curve analysis: 0.5°C/0.05s from 65°C to 95°C | Kolb et al. (2003) |
| | II646R | CGTGCCGCGCTCGACCATGYG | | | |
| ***16S*** | 16S-341F | CCTACGGGNGGCWGCAG | 464 | Denaturation: 95°C/3min<br>- 39 cycles: 95°C/15s; 60°C/1min; 80°C/5s (data acquisition)<br>- Melting curve analysis: 0.5°C/0.05s from 65°C to 95°C | Klindworth et al. (2012) |
| | 16S-805R | GACTACHVGGGTATCTAATCC | | | |

**Table S2** Summary of significant mixed linear models explaining methane ($CH_4$) fluxes across different variable subsets. Models within each subset are ranked and grouped according to fixed or random effects: site, sediment type (bare vs. seagrass), and subsite. The top-performing models within each group (i.e., those with the lowest AICc) are highlighted in bold. All models use $CH_4$ flux as the response variable, with predictors drawn from physical (A), gas-related (B), chemical (C), microbial (D-L), or combined (M) variable categories. Only models containing significant ($p < 0.05$) or marginally significant (# $p < 0.1$) predictors within each subset are presented. Variables included in each subset are listed at the top of their respective tables, n = 83.

#### A. Physical

| Subset of physical variables: Clay, Sand, Immersion_time, Hydro_forcing, Bathymetry, DBD, Moisture | | | | | |
|---|---|---|---|---|---|
| **Model** | **Formula: Fixed part** | **Random part** | **AICc (gls)** | **Fixed $R^2$** | **Global $R^2$** |
| **1** | **$log_{10}(CH_4\_flux)$ = 0.21 x Sand** | **~1\|Type** | **200** | **0.03** | **0.67** |
| **2** | **$log_{10}(CH_4\_flux)$ = - 0.50 x DBD + 0.39 x Sand - 0.37 x $log_{10}$(Moisture)** | **~1\|Type** | **200** | **0.06** | **0.73** |
| **3** | **$log_{10}(CH_4\_flux)$ = - 0.20 x $log_{10}$(Moisture)** | **~1\|Type** | **201** | **0.02** | **0.67** |
| **4** | **$log_{10}(CH_4\_flux)$ = - 0.20 x Clay** | **~1\|Type** | **201** | **0.02** | **0.67** |
| **5** | **$log_{10}(CH_4\_flux)$ = - 1.74 x Hydro_forcing + 0.94 x Bathymetry - 0.78 x DBD** | **~1+DBD\|Site** | **202** | **0.50** | **0.88** |
| 6 | $log_{10}(CH_4\_flux)$ = - 1.87 x Hydro_forcing + 1.51 x Immersion_time - 0.48 x DBD | ~1\|Site | 220 | 0.45 | 0.88 |
| 7 | $log_{10}(CH_4\_flux)$ = - 1.76 x Hydro_forcing + 1.12 x Bathymetry - 0.59 x DBD | ~1\|Site | 223 | 0.45 | 0.86 |
| **8** | **$log_{10}(CH_4\_flux)$ = - 0.57 x Hydro_forcing - 0.46 x DBD** | | **224 (232)** | **0.18** | |

### B. Gas-related

| Subset of Gas variables: $CO_2$_flux, $CO_2$_aerobic, $CH_4$_prod, $CO_2$_anaerobic, $CH_4$_oxi | | | | | |
|---|---|---|---|---|---|
| **Model** | **Formula: Fixed part** | **Random part** | **AICc (gls)** | **Fixed $R^2$** | **Global $R^2$** |
| **1** | **$log_{10}$($CH_4$_flux) = 0.35 x $CO_2$_flux** | **~1\|Subsite** | **190** | **0.11** | **0.69** |
| **2** | **$log_{10}$($CH_4$_flux) = 0.25 x $CO_2$_flux** | **~1\|Type** | **196** | **0.05** | **0.64** |
| **3** | **$log_{10}$($CH_4$_flux) = 0.44 x $log_{10}$($CO_2$_anaerobicobic) + 0.34 x $CO_2$_flux** | | **217 (225)** | **0.25** | |
| **4** | **$log_{10}$($CH_4$__flux) = 0.44 x $log_{10}$($CO_2$_anaerobic) + 0.34 x $CO_2$_flux** | **~1\|Site** | **228** | **0.27** | **0.28** |

### C. Chemical

| Subset of chemical variables: pH, Conductivity, N, $NH_4^+$, $NO_2^-$, $NO_3^-$, HI, $C_{min}$, $C_{org}$, CAR, SAR, MAR | | | | | |
|---|---|---|---|---|---|
| **Model** | **Formula: Fixed part** | **Random part** | **AICc (gls)** | **Fixed $R^2$** | **Global $R^2$** |
| **1** | **$log_{10}(CH_4$_flux) = 0.48 x CAR** | **~1\|Subsite** | **192** | **0.22** | **0.65** |
| 2 | $log_{10}(CH_4$_flux) = 0.53 x CAR - 0.18 x HI# | ~1\|Subsite | 193 | 0.25 | 0.66 |
| **3** | **$log_{10}(CH_4$_flux) = 0.27 x CAR - 0.22 x HI** | **~1\|Type** | **199** | **0.07** | **0.59** |
| **4** | **$log_{10}(CH_4$_flux) = - 0.26 x HI + 0.24 x MAR** | **~1\|Type** | **200** | **0.05** | **0.66** |
| **5** | **$log_{10}(CH_4$_flux) = 1.30 x CAR - 0.72 x SAR** | **~1+CAR\|Site** | **200** | **0.28** | **0.80** |
| 6 | $log_{10}(CH_4$_flux) = 0.21 x CAR | ~1\|Type | 201 | 0.04 | 0.55 |
| **7** | **$log_{10}(CH_4$_flux) = - 0.24 x HI + 0.22 x SAR** | **~1\|Type** | **201** | **0.05** | **0.62** |
| 8 | $log_{10}(CH_4$_flux) = 0.77 x CAR - 0.43 x MAR | ~1+CAR\|Site | 202 | 0.13 | 0.82 |
| 9 | $log_{10}(CH_4$_flux) = 0.66 x CAR + 0.54 x $C_{org}$ | ~1\|Site | 216 | 0.44 | 0.70 |
| 10 | $log_{10}(CH_4$_flux) = 1.18 x CAR - 0.60 x SAR | ~1\|Site | 223 | 0.38 | 0.49 |
| 11 | $log_{10}(CH_4$_flux) = 1.26 x CAR - 0.62 x SAR - 0.24 x HI | ~1\|Site | 223 | 0.41 | 0.54 |
| 12 | $log_{10}(CH_4$_flux) = 0.87 x CAR - 0.41 x MAR | ~1\|Site | 224 | 0.34 | 0.42 |
| **13** | **$log_{10}(CH_4$_flux) = 0.72 x CAR - 0.32 x MAR** | | **216 (224)** | **0.26** | |
| **14** | **$log_{10}(CH_4$_flux) = 0.84 x CAR - 0.40 x SAR#** | | **217 (224)** | **0.25** | |
| **15** | **$log_{10}(CH_4$_flux) = 0.54 x CAR - 0.22 x HI** | | **216 (225)** | **0.26** | |

### D. Microbial

| Subset of microbial variables: 16S, *mmoX*, *pmoA1a*, *pmoA1b*, *pmoAII*, *pmoAtot*, *mcrA*, DNA, pmoA_R, pmoA_D, mmoX_R, mmoX_D, mcrA_R, mcrA_D | | | | | |
|---|---|---|---|---|---|
| **Model** | **Formula: Fixed part** | **Random part** | **AICc (gls)** | **Fixed $R^2$** | **Global $R^2$** |
| **1** | **$log_{10}$($CH_4$_flux) = 0.25 x sqrt(pmoA_R) - 0.24 x mmoX_D - 0.22 x $log_{10}$(DNA)** | **~1\|Type** | **193** | **0.09** | **0.76** |
| 2 | $log_{10}$($CH_4$_flux) = 0.30 x sqrt(pmoA_R) - 0.28 x mmoX_D | ~1\|Type | 194 | 0.07 | 0.69 |
| 3 | $log_{10}$($CH_4$_flux) = - 0.30 x $log_{10}$(DNA) | ~1\|Type | 195 | 0.05 | 0.73 |
| 4 | $log_{10}$($CH_4$_flux) = 0.29 x sqrt(pmoA_R) - 0.24 x mmoX_D + 0.17 x $log_{10}$(mcrA_D) | ~1\|Type | 195 | 0.09 | 0.71 |
| **5** | **$log_{10}$($CH_4$_flux) = 0.41 x sqrt(pmoA_R)** | | **226 (231)** | **0.16** | |
| 6 | $log_{10}$($CH_4$_flux) = 0.41 x sqrt(pmoA_R) + 0.18 x $log_{10}$(mcrA_D)# | | 225 (233) | 0.18 | |
| **7** | **$log_{10}$($CH_4$_flux) = 0.41 x sqrt(pmoA_R)** | **~1\|Site** | **234** | **0.16** | **0.16** |
| 8 | $log_{10}$($CH_4$_flux) = 0.40 x sqrt(pmoA_D) + 0.25 x $log_{10}$(mcrA_D) | | 226 (235) | 0.16 | |
| 9 | $log_{10}$($CH_4$_flux) = 0.41 x sqrt(pmoA_R) + 0.18 x $log_{10}$(mcrA_D)# | ~1\|Site | 235 | 0.19 | 0.19 |

### E. Microbial qPCR

| Subset of microbial qPCR variables: *16S*, *mmoX*, *pmoA1a*, *pmoA1b*, *pmoAII*, *pmoAtot*, *mcrA*, DNA | | | | | |
|---|---|---|---|---|---|
| **Model** | **Formula: Fixed part** | **Random part** | **AICc (gls)** | **Fixed $R^2$** | **Global $R^2$** |
| **1** | **$log_{10}(CH_4$_flux) = - 0.30 x $log_{10}$(DNA)** | **~1\|Type** | **195** | **0.05** | **0.73** |
| **2** | **$log_{10}(CH_4$_flux) = - 0.15 x $log_{10}$(*pmoA1a*) + 0.12 x $log_{10}$(*pmoAtot*) - 0.11 x $log_{10}$(*pmoA1b*)** | | **238 (238)** | **0.08** | |
| **3** | **$log_{10}(CH_4$_flux) = 12.14 x $log_{10}$(*pmoAtot*) - 10.65 x $log_{10}$(*pmoA1b*) - 1.53 x $log_{10}$(*pmoA1a*)** | **~1\|Site** | **241** | **0.09** | **0.09** |
| 4 | $log_{10}(CH_4$_flux) = 0.77 x $log_{10}$(*pmoAII*) - 0.73 x $log_{10}$(*pmoA1a*) | | 236 (242) | 0.06 | |
| 5 | $log_{10}(CH_4$_flux) = 0.57 x $log_{10}$(*pmoAII*) - 0.55 x $log_{10}$(*mmoX*) | | 236 (243) | 0.06 | |
| 6 | $log_{10}(CH_4$_flux) = 0.49 x $log_{10}$(*16S*) - 0.48 x $log_{10}$(*mmoX*) | | 238 (244) | 0.07 | |

### F. Microbial diversity

| Subset of microbial qPCR diversity: pmoA_R, pmoA_D, mmoX_R, mmoX_D ,mcrA_R, mcrA_D | | | | | |
|---|---|---|---|---|---|
| **Model** | **Formula: Fixed part** | **Random part** | **AICc (gls)** | **Fixed $R^2$** | **Global $R^2$** |
| **1** | **$\log_{10}$($CH_4$_flux) = 0.30 x sqrt(pmoA_R) - 0.28 x mmoX_D** | **~1\|Type** | **194** | **0.07** | **0.70** |
| 2 | $\log_{10}$($CH_4$_flux) = - 0.28 x mmoX_D + 0.30 x sqrt(pmoA_R) + 0.17 x $\log_{10}$(mcrA_D) | ~1\|Type | 195 | 0.09 | 0.71 |
| **3** | **$\log_{10}$($CH_4$_flux) = 0.41 x sqrt(pmoA_R)** | | **226 (231)** | **0.16** | |
| 4 | $\log_{10}$($CH_4$_flux) = 0.41 x sqrt(pmoA_R) + 0.18 x $\log_{10}$(mcrA_D)# | | 225 (233) | 0.16 | |
| **5** | **$\log_{10}$($CH_4$_flux) = 0.41 x sqrt(pmoA_R)** | **~1\|Site** | **234** | **0.16** | **0.16** |
| 6 | $\log_{10}$($CH_4$_flux) = 0.41 x sqrt(pmoA_R) + 0.18 x $\log_{10}$(mcrA_D) | ~1\|Site | 235 | 0.19 | 0.19 |
| 7 | $\log_{10}$($CH_4$_flux) = 0.40 x sqrt(pmoA_D) + 0.25 x $\log_{10}$(mcrA_D) | | 226 (235) | 0.16 | |

### G. mcrA

| Subset of microbial mcrA variables: *16S*, *mcrA*, DNA, mcrA_R, mcrA_D | | | | | |
|---|---|---|---|---|---|
| **Model** | **Formula: Fixed part** | **Random part** | **AICc (gls)** | **Fixed $R^2$** | **Global $R^2$** |
| **1** | **$\log_{10}$($CH_4$_flux) = - 0.30 x $\log_{10}$(DNA)** | **~1\|Type** | **194** | **0.05** | **0.73** |
| **2** | **$\log_{10}$($CH_4$_flux) = - 0.27 x $\log_{10}$(DNA) + 0.16 x $\log_{10}$(mcrA_D)** | **~1\|Type** | **196** | **0.06** | **0.74** |

#### H. mmoX

| Subset of microbial mmoX variables: *16S*, *mmoX*, DNA, mmoX_R, mmoX_D | | | | | |
|---|---|---|---|---|---|
| **Model** | **Formula: Fixed part** | **Random part** | **AICc (gls)** | **Fixed $R^2$** | **Global $R^2$** |
| **1** | **log$_{10}$(CH$_4$_flux) = - 0.30 x log$_{10}$(DNA)** | **~1\|Type** | **194** | **0.05** | **0.73** |
| **2** | **log$_{10}$(CH$_4$_flux) = 0.49 x log$_{10}$(*16S*) - 0.48 x log$_{10}$(*mmoX*)** | | **238 (244)** | **0.04** | |
| **3** | **log$_{10}$(CH$_4$_flux) = 0.51 x log$_{10}$(*16S*) - 0.50 x log$_{10}$(*mmoX*)** | **~1\|Site** | **246** | **0.07** | **0.09** |

#### I. pmoA

| Subset of microbial pmoA variables: *16S*, *pmoA1a*, *pmoA1b*, *pmoAII*, *pmoAtot*, DNA, pmoA_R, pmoA_D | | | | | |
|---|---|---|---|---|---|
| **Model** | **Formula: Fixed part** | **Random part** | **AICc (gls)** | **Fixed $R^2$** | **Global $R^2$** |
| **1** | **log$_{10}$(CH$_4$_flux) = - 0.30 x log$_{10}$(DNA)** | **~1\|Type** | **194** | **0.05** | **0.73** |
| **2** | **log$_{10}$(CH$_4$_flux) = - 0.26 x log$_{10}$(DNA) + 0.18 x log$_{10}$(pmoA_R)** | **~1\|Type** | **195** | **0.07** | **0.71** |
| **3** | **log$_{10}$(CH$_4$_flux) = 0.41 x sqrt(pmoA_R)** | | **226 (231)** | **0.16** | |
| **4** | **log$_{10}$(CH$_4$_flux) = 0.41 x sqrt(pmoA_R)** | **~1\|Site** | **234** | **0.16** | **0.16** |
| 5 | log$_{10}$(CH$_4$_flux) = 10.11 x log$_{10}$(*pmoAtot*) - 9.30 x log$_{10}$(*pmoA1b*) - 1.00 x log$_{10}$(*pmoA1a*)# + 0.39 x sqrt(pmoA_D) | ~1\|Site | 235 | 0.19 | 0.20 |
| 6 | log$_{10}$(CH$_4$_flux) = 9.29 x log$_{10}$(*pmoAtot*) - 8.86 x log$_{10}$(*pmoA1b*) - 1.07 x log$_{10}$(*pmoA1a*)# + 0.48 x log$_{10}$(*16S*)# + 0.40 x sqrt(pmoA_D) | ~1\|Site | 236 | 0.28 | 0.22 |

**J. mcrA genus**

| Subset of mcrA genus variables: *Methermicoccus, Methanococcoides, Methanolobus, Methanogenium, Methanosarcina, Methanobacterium, Methanoculleus, Methanocalculus, Methanolinea, Methanopyrus, Methanocaldococcus, Methanococcus_fervidus, Methanolubus, Methanohalophilus* | | | | | |
|---|---|---|---|---|---|
| **Model** | **Formula: Fixed part** | **Random part** | **AICc (gls)** | **Fixed $R^2$** | **Global $R^2$** |
| **1** | **$\log_{10}(CH_4\_flux)$ = 0.21 x *Methanopyrus*** | **~1\|Type** | **200** | **0.03** | **0.65** |
| **2** | **$\log_{10}(CH_4\_flux)$ = 0.17 x *Methanocaldococcus*** | **~1\|Type** | **202** | **0.02** | **0.61** |
| **3** | **$\log_{10}(CH_4\_flux)$ = 0.26 x *Methanolobus* + 0.25 x *Methanocaldococcus*** | | **233 (241)** | **0.09** | |
| 4 | $\log_{10}(CH_4\_flux)$ = 0.21 x *Methanocaldococcus* | | 237 (242) | 0.03 | |
| 5 | $\log_{10}(CH_4\_flux)$ = 0.25 x *Methanolobus* + 0.24 x *Methanocaldococcus* - 0.19 x *Methanolinea*# | | 232 (243) | 0.12 | |
| **6** | **$\log_{10}(CH_4\_flux)$ = 0.22 x *Methanocaldococcus*** | **~1\|Site** | **244** | **0.05** | **0.05** |
| **7** | **$\log_{10}(CH_4\_flux)$ = - 0.24 x *Methanolinea*** | **~1\|Site** | **244** | **0.06** | **0.08** |
| **8** | **$\log_{10}(CH_4\_flux)$ = 0.23 x *Methanolobus*** | **~1\|Site** | **244** | **0.05** | **0.05** |

**K. mmoX genus**

Subset of mmoX genus variables: *Methylocella, Methyloferula, Methylocystis_m, Methylosinus_m, Methylomicrobium_m, Methylovulum_m, Methylomonas_m, Methylococcus_m, Methylomagnum_m, Methylocaldum_m, Pseudomonas, Mycobacterium, Rhodococcus, Methylocella_prmA, Methylibium, Bradyrhizobium, Paracoccus, Gordonia*

| Model | Formula: Fixed part | Random part | AICc (gls) | Fixed $R^2$ | Global $R^2$ |
|---|---|---|---|---|---|
| **1** | **log$_{10}$($CH_4$_flux) = - 0.19 x *Methyloferula*** | **~1\|Subsite** | **193** | **0.03** | **0.68** |
| 2 | log$_{10}$($CH_4$_flux) = - 0.19 x *Methyloferula* + 0.13 x *Methylomicrobium_m*# | ~1\|Subsite | 195 | 0.05 | 0.69 |
| **3** | **log$_{10}$($CH_4$_flux) = 0.16 x *Methylocella*#** | **~1\|Type** | **202** | **0.02** | **0.62** |
| 4 | log$_{10}$($CH_4$_flux) = 0.27 x *Methylomicrobium_m* - 0.24 x *Methylomonas_m* + 0.20 x *Methylocella* - 0.16 x *Methyloferula* | ~1\|Type | 209 | 0.06 | 0.66 |
| **5** | **log$_{10}$($CH_4$_flux) = - 0.43 x *Methylomagnum_m* + 0.37 x *Methylocaldum_m*** | | **237 (244)** | **0.05** | |
| **6** | **log$_{10}$($CH_4$_flux) = 0.20 x *Methylomicrobium_m*#** | **~1\|Site** | **245** | **0.04** | **0.04** |
| **7** | **log$_{10}$($CH_4$_flux) = - 0.43 x *Methylomagnum_m* + 0.37 x *Methylocaldum_m*** | **~1\|Site** | **246** | **0.07** | **0.07** |
| 8 | log$_{10}$($CH_4$_flux) = - 0.41 x *Methylomagnum_m* + 0.32 x *Methylocaldum_m*# + 0.19 x *Methylomicrobium_m*# | | 236 (246) | 0.07 | |

#### L. pmoA genus

| Subset of pmoA genus variables: *Methylocaldum_p, Methylogaea, Methylosinus_p, Methylocystis_p, Methylocapsa, Methylomonas_p, Methylomagnum_p, Candidatus_Binatia, Candidatus_Methylomirabilis, Methylomarinum, Methylomicrobium_p, Methyloglobulus, Methylovulum_p, Methylosoma, Methylobacter, Methylococcus_p, Clonothrix* | | | | | |
|---|---|---|---|---|---|
| **Model** | **Formula: Fixed part** | **Random part** | **AICc (gls)** | **Fixed $R^2$** | **Global $R^2$** |
| **1** | **$log_{10}(CH_4$_flux) = 0.17 x *Methylogaea*** | **~1\|Subsite** | **194** | **0.03** | **0.69** |
| **2** | **$log_{10}(CH_4$_flux) = 0.16 x *Methyloglobulus#*** | **~1\|Type** | **203** | **0.02** | **0.60** |
| **3** | **$log_{10}(CH_4$_flux) = 0.15 x *Methylomarinum#*** | **~1\|Type** | **203** | **0.02** | **0.60** |
| **4** | **$log_{10}(CH_4$_flux) = 0.15 x *Methylogaea#*** | **~1\|Type** | **203** | **0.01** | **0.63** |
| 5 | $log_{10}(CH_4$_flux) = 0.16 x *Methyloglobulus* + 0.15 x *Methylomarinum#* | ~1\|Type | 205 | 0.04 | 0.60 |
| 6 | $log_{10}(CH_4$_flux) = 0.27 x *Methyloglobulus* + 0.24 x *Methylococcus_p* | | 232 (240) | 0.10 | |
| **7** | **$log_{10}(CH_4$_flux) = 0.28 x *Methyloglobulus* + 0.22 x *Methylococcus_p* + 0.21 x *Methylomarinum*** | | **230 (241)** | **0.14** | |
| 8 | $log_{10}(CH_4$_flux) = 0.27 x *Methyloglobulus* + 0.24 x *Methylococcus_p* | ~1\|Site | 242 | 0.12 | 0.12 |
| **9** | **$log_{10}(CH_4$_flux) = 0.26 x *Methyloglobulus*** | **~1\|Site** | **242** | **0.07** | **0.07** |
| 10 | $log_{10}(CH_4$_flux) = 0.28 x *Methyloglobulus* + 0.22 x *Methylococcus_p* + 0.21 x *Methylomarinum* | ~1\|Site | 243 | 0.16 | 0.16 |
| 11 | $log_{10}(CH_4$_flux) = 0.27 x *Methyloglobulus* + 0.23 x *Methylomarinum* | ~1\|Site | 243 | 0.12 | 0.12 |
| 12 | $log_{10}(CH_4$_flux) = 0.22 x *Methylococcus_p* | ~1\|Site | 244 | 0.05 | 0.05 |
| 13 | $log_{10}(CH_4$_flux) = 0.22 x *Methylomarinum* | ~1\|Site | 244 | 0.05 | 0.05 |

**M. Combined**

Subset of combined variables: $CO_2$_flux, $CO_2$_anaerobic, CAR, HI, MAR, SAR, DBD, Sand, Hydro_forcing, Bathymetry, pmoA_R, DNA, mmoX_D, *pmoAII*, *16S*, pmoA_D, *Methyloferula, Methylogaea, Methanopyrus, Methanocaldococcus, Methyloglobulus, Methylomarinum, Methylomicrobium_m, Methylomonas_m, Methylocella, Methylococcus_p, Methanolobus, Methanolinea, Methylomagnum_m, Methylocaldum_m*

| Model | Formula: Fixed part | Random part | AICc (gls) | Fixed $R^2$ | Global $R^2$ |
|---|---|---|---|---|---|
| 1 | $log_{10}(CH_4$_flux) = 0.96 x CAR - 0.56 x MAR + 0.41 x $CO_2$_flux - 0.18 x *Methyloferula* | ~1\|Subsite | 185 | 0.44 | 0.70 |
| **2** | **$log_{10}(CH_4$_flux) = 0.95 x CAR - 0.56 x MAR + 0.43 x $CO_2$_flux** | **~1\|Subsite** | **186** | **0.43** | **0.77** |
| **3** | **$log_{10}(CH_4$_flux) = 0.64 x CAR - 0.39 x MAR + 0.33 x $CO_2$_flux - 0.29 x $log_{10}$(DNA)** | **~1\|Type** | **186** | **0.21** | **0.65** |
| 4 | $log_{10}(CH_4$_flux) = 0.66 x CAR - 0.40 x MAR + 0.32 x $CO_2$_flux - 0.31 x $log_{10}$(DNA) - 0.15 x *Methyloferula* | ~1\|Type | 188 | 0.23 | 0.69 |
| **5** | **$log_{10}(CH_4$_flux) = 1.66 x CAR - 1.28 x Hydro_forcing - 1.27 x SAR + 0.77 x Bathymetry + 0.46 x $CO_2$_flux** | **~1\|Site** | **193** | **0.57** | **0.88** |
| 6 | $log_{10}(CH_4$_flux) = 1.10 x CAR - 0.77 x MAR + 0.52 x $CO_2$_flux + 0.22 x $log_{10}$(*16S*) | | 186 (201) | 0.50 | |
| **7** | **$log_{10}(CH_4$_flux) = 1.10 x CAR - 0.72 x MAR + 0.51 x $CO_2$_flux + 0.27 x $log_{10}$(*16S*) - 0.20 x HI** | | **183 (201)** | **0.53** | |
| **8** | **$log_{10}(CH_4$_flux) = 1.15 x CAR - 0.85 x MAR + 0.50 x $CO_2$_flux + 0.24 x $log_{10}$(*16S*) + 0.17 x *Methanopyrus*** | | **183 (202)** | **0.52** | |
| **9** | **$log_{10}(CH_4$_flux) = 1.12 x CAR - 0.87 x MAR + 0.50 x $CO_2$_flux + 0.24 x $log_{10}$(*16S*) - 0.18 x $log_{10}$(DNA)** | | **184 (202)** | **0.52** | |
| **10** | **$log_{10}(CH_4$_flux) = 1.07 x CAR - 0.74 x MAR + 0.50 x $CO_2$_flux + 0.24 x $log_{10}$(*16S*) + 0.16 x *Methanocaldococcus*** | | **184 (202)** | **0.52** | |
| **11** | **$log_{10}(CH_4$_flux) = 1.11 x CAR - 0.79 x MAR + 0.50 x $CO_2$_flux + 0.25 x $log_{10}$(*16S*) - 0.16 x *Methyloferula*** | | **184 (203)** | **0.52** | |
| 12 | $log_{10}(CH_4$_flux) = 0.99 x CAR - 0.60 x MAR + 0.50 x $CO_2$_flux | | 190 (202) | 0.47 | |
| **13** | **$log_{10}(CH_4$_flux) = 1.12 x CAR - 0.77 x MAR + 0.57 x $CO_2$_flux + 0.31 x $log_{10}$(*16S*) + 0.18 x Bathymetry#** | | **185 (203)** | **0.52** | |

**Table S3** Microbial characteristics of the studied sites. A DNA content and qPCR results. B Diversity indices. Values are expressed as mean ± standard error (SE), n = 83. Sites are ordered along a gradient from the most negative to the most positive values along PC1 (see Figure 2), which reflects a multivariate ordination of environmental and microbial descriptors. Differences among sites were assessed using non-parametric Kruskal–Wallis tests, followed by Dunn's post hoc tests with Benjamini–Hochberg correction for multiple comparisons. Different letters indicate significant differences among sites for each variable ($p < 0.05$); groups sharing at least one letter are not significantly different.

**A**

| **Site_ID** | **DNA** ng m⁻² | | ***16S*** copies m⁻² | | ***mcrA*** copies m⁻² | | ***mmoX*** copies m⁻² | | ***pmoA1a*** copies m⁻² | | ***pmoA1b*** copies m⁻² | | ***pmoAII*** copies m⁻² | | ***pmoAtot*** copies m⁻² | |
|---|---|---|---|---|---|---|---|---|---|---|---|---|---|---|---|---|
| Gail | 1.5e+09 ± 2.5e+08 | bc | 6.2e+13 ± 4.5e+12 | d | 1.3e+10 ± 3.6e+09 | b | 5.4e+09 ± 4.8e+08 | c | 2.9e+10 ± 2.3e+09 | c | 4.1e+11 ± 3.1e+10 | d | 2.3e+10 ± 2.8e+09 | c | 4.6e+11 ± 3.5e+10 | d |
| Garr | 8.6e+08 ± 2.1e+08 | ab | 6.7e+13 ± 1.3e+13 | d | 2.0e+10 ± 4.2e+09 | ab | 5.5e+09 ± 9.7e+08 | c | 3.1e+10 ± 5.1e+09 | c | 4.6e+11 ± 7.9e+10 | d | 3.3e+10 ± 9.7e+09 | c | 5.2e+11 ± 9.2e+10 | d |
| Jacq | 1.7e+09 ± 5.1e+08 | abc | 9.5e+13 ± 8.5e+12 | cd | 1.1e+10 ± 2.2e+09 | b | 7.2e+09 ± 6.4e+08 | bc | 4.0e+10 ± 3.0e+09 | bc | 5.9e+11 ± 4.2e+10 | cd | 4.1e+10 ± 3.9e+09 | bc | 6.7e+11 ± 4.8e+10 | cd |
| Germ | 1.5e+09 ± 1.7e+08 | c | 1.4e+14 ± 2.4e+13 | bcd | 1.6e+10 ± 3.2e+09 | ab | 1.3e+10 ± 1.8e+09 | ab | 7.7e+10 ± 1.2e+10 | ab | 9.3e+11 ± 1.3e+11 | ac | 7.2e+10 ± 1.1e+10 | ab | 1.1e+12 ± 1.5e+11 | ac |
| Comp | 8.9e+08 ± 1.5e+08 | abc | 1.7e+14 ± 1.5e+13 | abc | 1.4e+10 ± 4.3e+09 | b | 1.2e+10 ± 1.2e+09 | ab | 7.1e+10 ± 6.9e+09 | ab | 8.5e+11 ± 7.9e+10 | ac | 7.0e+10 ± 9.4e+09 | ab | 9.9e+11 ± 9.4e+10 | ac |
| Ande | 1.0e+09 ± 1.1e+08 | abc | 1.9e+14 ± 2.1e+13 | ab | 3.8e+10 ± 7.9e+09 | a | 2.1e+10 ± 2.4e+09 | a | 8.5e+10 ± 6.4e+09 | a | 1.4e+12 ± 1.0e+11 | ab | 1.0e+11 ± 8.9e+09 | a | 1.6e+12 ± 1.1e+11 | ab |
| Font | 6.6e+08 ± 5.5e+07 | a | 3.1e+14 ± 3.5e+13 | a | 3.4e+10 ± 6.1e+09 | a | 2.4e+10 ± 3.3e+09 | a | 1.2e+11 ± 1.5e+10 | a | 1.8e+12 ± 1.8e+11 | b | 1.1e+11 ± 1.6e+10 | a | 2.0e+12 ± 2.0e+11 | b |
| All | 1.2e+09 ± 1.0e+08 | | 1.5e+14 ± 1.1e+13 | | 2.1e+10 ± 2.1e+09 | | 1.2e+10 ± 1.0e+09 | | 6.5e+10 ± 4.7e+09 | | 9.2e+11 ± 6.4e+10 | | 6.5e+10 ± 5.0e+09 | | 1.0e+12 ± 7.3e+10 | |

**B**

| Site_ID | mcrA_R | | mcrA_D | | mmoX_R | | mmoX_D | | pmoA_R | | pmoA_D | |
|---|---|---|---|---|---|---|---|---|---|---|---|---|
| Gail | 50.2 ± 8.3 | bc | 3.1 ± 0.1 | ab | 27.1 ± 1.5 | b | 2.9 ± 0.1 | ab | 5.4 ± 0.8 | c | 0.5 ± 0.1 | b |
| Garr | 53.5 ± 3.6 | c | 3.5 ± 0.1 | b | 28.6 ± 1.7 | ab | 2.7 ± 0.1 | b | 8.1 ± 0.9 | bc | 0.4 ± 0.1 | b |
| Jacq | 39.1 ± 10.5 | a | 2.9 ± 0.2 | a | 28.0 ± 1.6 | b | 2.7 ± 0.1 | b | 6.8 ± 0.8 | bc | 0.4 ± 0.1 | b |
| Germ | 24.9 ± 2.0 | a | 2.7 ± 0.1 | a | 29.8 ± 1.7 | ab | 3.0 ± 0.0 | ab | 7.2 ± 0.5 | bc | 0.6 ± 0.1 | ab |
| Comp | 26.6 ± 2.1 | a | 2.8 ± 0.1 | a | 35.4 ± 1.3 | a | 3.1 ± 0.0 | a | 9.5 ± 0.8 | ab | 0.5 ± 0.1 | b |
| Ande | 31.0 ± 2.8 | ab | 2.8 ± 0.1 | a | 30.4 ± 1.5 | ab | 3.0 ± 0.0 | ab | 14.5 ± 1.5 | a | 1.1 ± 0.1 | a |
| Font | 27.8 ± 3.1 | a | 2.7 ± 0.1 | a | 24.8 ± 2.0 | b | 2.8 ± 0.1 | b | 10.2 ± 2.1 | abc | 0.8 ± 0.2 | ab |
| All | 36.3 ± 2.4 | | 2.9 ± 0.0 | | 29.1 ± 0.7 | | 2.9 ± 0.0 | | 8.8 ± 0.5 | | 0.6 ± 0.0 | |

**Table S4** Characteristics of the studied sites under bare and vegetated (seagrass) conditions, with variables grouped into general and physical features (A–B) and chemical variables (C–D). Significant differences (Wilcoxon tests) between bare and vegetated sediments are indicated as follows: # ($p < 0.1$), * ($p < 0.05$), ** ($p < 0.01$) ), *** ($p < 0.001$), and NS (not significant), n = 83.

**A**

| Site | Site_ID | Type (Subsite) | Longitude °E. WEGS84 | Latitude °N. WGS84 | Plant_cover | | Sediment texture | Clay % | | Silt % | | Sand % | |
|---|---|---|---|---|---|---|---|---|---|---|---|---|---|
| Comprian | Comp | Bare | -1.070611 | 44.679473 | 0 | | Silty sand | 3.0 | | 47.4 | | 49.6 | |
| | | Seagrass | -1.072061 | 44.679063 | 0.9 | | Silty sand | 3.8 | | 47.8 | | 48.4 | |
| Jacquets | Jacq | Bare | -1.18081 | 44.72352 | 0 | | Silty sand | 1.8 | | 29.5 | | 68.7 | |
| | | Seagrass | -1.18128 | 44.72466 | 0.9 | | Silty sand | 1.8 | | 45.2 | | 53.0 | |
| Germanan | Germ | Bare | -1.13384771347 | 44.7116661072 | 0 | | Sandy silt | 4.1 | | 63.4 | | 32.5 | |
| | | Seagrass | -1.13384771347 | 44.7116661072 | 0.9 | | Sandy silt | 4.1 | | 55.4 | | 40.5 | |
| Fontaines | Font | Bare | -1.07578 | 44.71885 | 0 | | Sand | 0.6 | | 8.4 | | 91.0 | |
| | | Seagrass | -1.0754 | 44.71874 | 0.9 | | Sandy silt | 3.6 | | 52.4 | | 44.0 | |
| Garrèche | Garr | Bare | -1.1205807 | 44.7047449 | 0 | | Silty sand | 1.7 | | 26 | | 72.3 | |
| | | Seagrass | -1.1208349 | 44.7040884 | 0.4 | | Silty sand | 2.5 | | 37.8 | | 59.7 | |
| Gaillard | Gail | Bare | -1.0994219 | 44.6627752 | 0 | | Sandy silt | 3.5 | | 61.2 | | 35.3 | |
| | | Seagrass | -1.09981 | 44.66241 | 0.9 | | Sandy silt | 3.2 | | 60.4 | | 36.4 | |
| Andernos | Ande | Bare | -1.12038 | 44.74512 | 0 | | Sand | 1.3 | | 23.7 | | 75.0 | |
| | | Seagrass | -1.12138 | 44.74513 | 0.9 | | Sand | 1.3 | | 21.5 | | 77.2 | |
| All sites | All | Bare | NA | NA | 0.0 | *** | NA | 2.3 ± 0.2 | * | 37.1 ± 3.2 | NS | 60.9 ± 3.2 | NS |
| | | Seagrass | | | 0.8 ± 0.1 | | | 2.9 ± 0.2 | | 45.8 ± 2.0 | | 51.3 ± 2.0 | |

**B**

| Site | Site_ID | Type (Subsite) | Bathymetry m | | Hydrodynamic forcing | | Immersion_time h $day^{-1}$ | | DBD g $cm^{-3}$ | | Moisture % | |
|---|---|---|---|---|---|---|---|---|---|---|---|---|
| Comprian | Comp | Bare | 1.4 | | 2.1e+05 | | 18.3 | | 1.1 | | 39.3 ± 3.0 | ** |
| | | Seagrass | -0.2 | | 8.3e+04 | | 10.6 | | 1.0 | | 55.0 ± 3.4 | |
| Jacquets | Jacq | Bare | 2.1 | | 2.0e+05 | | 18.1 | | 1.2 | | 36.3 ± 3.8 | ** |
| | | Seagrass | 0.6 | | 1.4e+05 | | 12.2 | | 0.9 | | 80.4 ± 5.0 | |
| Germanan | Germ | Bare | 1.7 | | 1.3e+05 | | 18.0 | | 0.8 | | 62.8 ± 3.5 | NS |
| | | Seagrass | 1.7 | | 1.3e+05 | | 18.0 | | 1.2 | | 58.1 ± 4.5 | |
| Fontaines | Font | Bare | -1.1 | | 0 | | 1.7 | | 1.7 | | 23.2 ± 0.5 | ** |
| | | Seagrass | -1.1 | | 0 | | 1.7 | | 1.2 | | 38.5 ± 2.3 | |
| Garrèche | Garr | Bare | 1.2 | | 2.3e+05 | | 17.3 | | 1.1 | | 52.7 ± 5.3 | NS |
| | | Seagrass | 1.1 | | 1.1e+05 | | 17.3 | | 1.1 | | 56.7 ± 7.3 | |
| Gaillard | Gail | Bare | 0.3 | | 3.9e+05 | | 13.1 | | 0.8 | | 79.8 ± 3.6 | NS |
| | | Seagrass | 0.5 | | 2.8e+05 | | 13.1 | | 0.8 | | 74.0 ± 5.5 | |
| Andernos | Ande | Bare | -0.8 | | 0 | | 1.7 | | 1.4 | | 34.8 ± 1.3 | ** |
| | | Seagrass | -0.5 | | 0 | | 5.6 | | 1.1 | | 52.1 ± 2.7 | |
| All sites | All | Bare | 0.7 ± 0.2 | # | 1.6e+05 ± 2.0e+04 | * | 12.5 ± 1.1 | NS | 1.2 ± 0.0 | # | 47.2 ± 3.1 | ** |
| | | Seagrass | 0.3 ± 0.1 | | 1.1e+05 ± 1.4e+04 | | 11.2 ± 0.9 | | 1.0 ± 0.0 | | 59.2 ± 2.6 | |

**C**

| Site_ID | Type (Subsite) | pH | | Conductivity mS $cm^{-1}$ | | N % | | $NH_4^+$ µg N-$NH_4$ $g^{-1}$ dry soil | | $NO_2^-$ µg N-$NO_2$ $g^{-1}$ dry soil | | $NO_3^-$ µg N-$NO_3$ $g^{-1}$ dry soil | |
|---|---|---|---|---|---|---|---|---|---|---|---|---|---|
| Comp | Bare | 8.5 ± 0.0 | NS | 3.5 ± 0.3 | NS | 0.070 ± 0.008 | ** | 19.7 ± 5.3 | NS | 0.020 ± 0.003 | NS | 0.450 ± 0.213 | NS |
| | Seagrass | 8.5 ± 0.0 | | 3.2 ± 0.2 | | 0.119 ± 0.011 | | 10.6 ± 2.1 | | 0.019 ± 0.001 | | 0.587 ± 0.088 | |
| Jacq | Bare | 8.6 ± 0.1 | * | 4.3 ± 0.4 | * | 0.068 ± 0.010 | ** | 6.4 ± 0.6 | * | 0.017 ± 0.001 | NS | 0.409 ± 0.095 | NS |
| | Seagrass | 8.3 ± 0.1 | | 6.9 ± 0.8 | | 0.169 ± 0.007 | | 9.7 ± 0.9 | | 0.014 ± 0.001 | | 0.534 ± 0.140 | |
| Germ | Bare | 7.3 ± 0.6 | NS | 6.6 ± 0.5 | NS | 0.139 ± 0.009 | NS | 6.6 ± 1.0 | # | 0.014 ± 0.003 | NS | 0.105 ± 0.101 | # |
| | Seagrass | 8.1 ± 0.2 | | 6.0 ± 0.8 | | 0.129 ± 0.014 | | 8.1 ± 0.5 | | 0.015 ± 0.002 | | 0.439 ± 0.122 | |
| Font | Bare | 6.8 ± 0.1 | ** | 4.4 ± 1.2 | # | 0.024 ± 0.008 | ** | 4.3 ± 0.8 | # | 0.009 ± 0.001 | ** | 0.219 ± 0.031 | NS |
| | Seagrass | 8.4 ± 0.1 | | 4.3 ± 0.3 | | 0.067 ± 0.011 | | 6.8 ± 1.3 | | 0.017 ± 0.002 | | 0.163 ± 0.017 | |
| Garr | Bare | 8.4 ± 0.1 | NS | 5.9 ± 0.6 | NS | 0.116 ± 0.009 | NS | 5.9 ± 0.4 | NS | 0.020 ± 0.002 | NS | 0.253 ± 0.160 | NS |
| | Seagrass | 8.4 ± 0.1 | | 6.4 ± 0.6 | | 0.118 ± 0.009 | | 6.3 ± 0.7 | | 0.021 ± 0.010 | | 0.431 ± 0.208 | |
| Gail | Bare | 8.0 ± 0.0 | NS | 7.4 ± 1.0 | NS | 0.202 ± 0.009 | # | 2.5 ± 0.4 | # | 0.016 ± 0.001 | NS | 0.072 ± 0.014 | NS |
| | Seagrass | 7.4 ± 0.4 | | 6.6 ± 1.0 | | 0.158 ± 0.017 | | 5.0 ± 0.8 | | 0.018 ± 0.001 | | 0.152 ± 0.081 | |
| Ande | Bare | 6.8 ± 0.5 | * | 4.0 ± 0.2 | ** | 0.155 ± 0.084 | # | 2.8 ± 0.8 | * | 0.020 ± 0.009 | NS | 0.064 ± 0.016 | NS |
| | Seagrass | 7.8 ± 0.1 | | 6.2 ± 0.4 | | 0.107 ± 0.015 | | 5.4 ± 1.1 | | 0.017 ± 0.001 | | 0.082 ± 0.010 | |
| All | Bare | 7.8 ± 0.2 | NS | 5.2 ± 0.3 | NS | 0.111 ± 0.015 | * | 6.6 ± 1.0 | ** | 0.016 ± 0.002 | NS | 0.219 ± 0.044 | * |
| | Seagrass | 8.1 ± 0.1 | | 5.7 ± 0.3 | | 0.124 ± 0.007 | | 7.4 ± 0.5 | | 0.017 ± 0.001 | | 0.341 ± 0.050 | |

**D**

| Site_ID | Type (Subsite) | $C_{mineral}$ % | | $C_{organic}$ % | | HI | | SAR cm $y^{-1}$ | | MAR g $cm^{-2}$ $y^{-1}$ | | CAR g $CO_2$-eq $m^{-2}$ $y^{-1}$ | | Climate regulation g $CO_2$-eq $m^{-2}$ $y^{-1}$ | |
|---|---|---|---|---|---|---|---|---|---|---|---|---|---|---|---|
| Comp | Bare | 0.082 ± 0.018 | * | 1.1 | | 190.6 ± 3.0 | NS | 0.09 | | 0.10 | | 33.8 | | 21.6 ± 2.8 | ** |
| | Seagrass | 0.360 ± 0.095 | | 0.6 | | 201.0 ± 5.2 | | 0.09 | | 0.06 | | 12.6 | | 5.1 ± 1.1 | |
| Jacq | Bare | 0.580 ± 0.055 | * | 1.7 | | 210.3 ± 6.7 | * | 0.23 | | 0.33 | | 64.4 | | 62.3 ± 0.2 | ** |
| | Seagrass | 0.267 ± 0.065 | | 1.4 | | 238.8 ± 6.0 | | 0.45 | | 0.44 | | 164.8 | | 85.3 ± 3.1 | |
| Germ | Bare | 0.158 ± 0.089 | NS | 1.5 | | 208.2 ± 17.4 | ** | 0.28 | | 0.13 | | 38.8 | | 48.5 ± 1.7 | ** |
| | Seagrass | 0.067 ± 0.004 | | 1.5 | | 246.7 ± 3.6 | | 0.11 | | 0.09 | | 22.9 | | 18.6 ± 1.6 | |
| Font | Bare | 0.020 ± 0.004 | * | 0.1 | | 243.0 ± 10.6 | NS | 0.23 | | 0.37 | | 21.7 | | 2.3 ± 2.2 | ** |
| | Seagrass | 0.096 ± 0.022 | | 0.8 | | 240.5 ± 1.4 | | 0.26 | | 0.30 | | 78.9 | | 53.9 ± 4.4 | |
| Garr | Bare | 0.751 ± 0.090 | NS | 1.3 | | 188.3 ± 7.8 | NS | 0 | | 0 | | 0 | | -13.0 ± 5.1 | ** |
| | Seagrass | 0.646 ± 0.184 | | 0.6 | | 201.8 ± 3.6 | | 0.32 | | 0.33 | | 92.2 | | 78.9 ± 3.0 | |
| Gail | Bare | 0.164 ± 0.015 | * | 0.9 | | 229.0 ± 2.2 | NS | 0 | | 0 | | 0 | | 5.2 ± 1.8 | ** |
| | Seagrass | 0.107 ± 0.019 | | 2.0 | | 209.3 ± 13.1 | | 0.15 | | 0.11 | | 62.6 | | 47.8 ± 1.7 | |
| Ande | Bare | 0.011 ± 0.003 | ** | 0.5 | | 250.7 ± 12.3 | # | 0.29 | | 0.40 | | 74.5 | | 55.8 ± 5.3 | ** |
| | Seagrass | 0.081 ± 0.029 | | 1.1 | | 242.8 ± 4.4 | | 0.35 | | 0.28 | | 129.9 | | 104.0 ± 1.6 | |
| All | Bare | 0.256 ± 0.047 | NS | 1.0 ± 0.1 | NS | 217.8 ± 5.0 | NS | 0.162 ± 0.019 | ** | 0.192 ± 0.026 | NS | 33.3 ± 4.3 | *** | 26.2 ± 4.5 | *** |
| | Seagrass | 0.232 ± 0.042 | | 1.1 ± 0.1 | | 225.9 ± 3.7 | | 0.247 ± 0.020 | | 0.230 ± 0.021 | | 80.6 ± 7.9 | | 56.2 ± 5.3 | |

### Supplementary figures

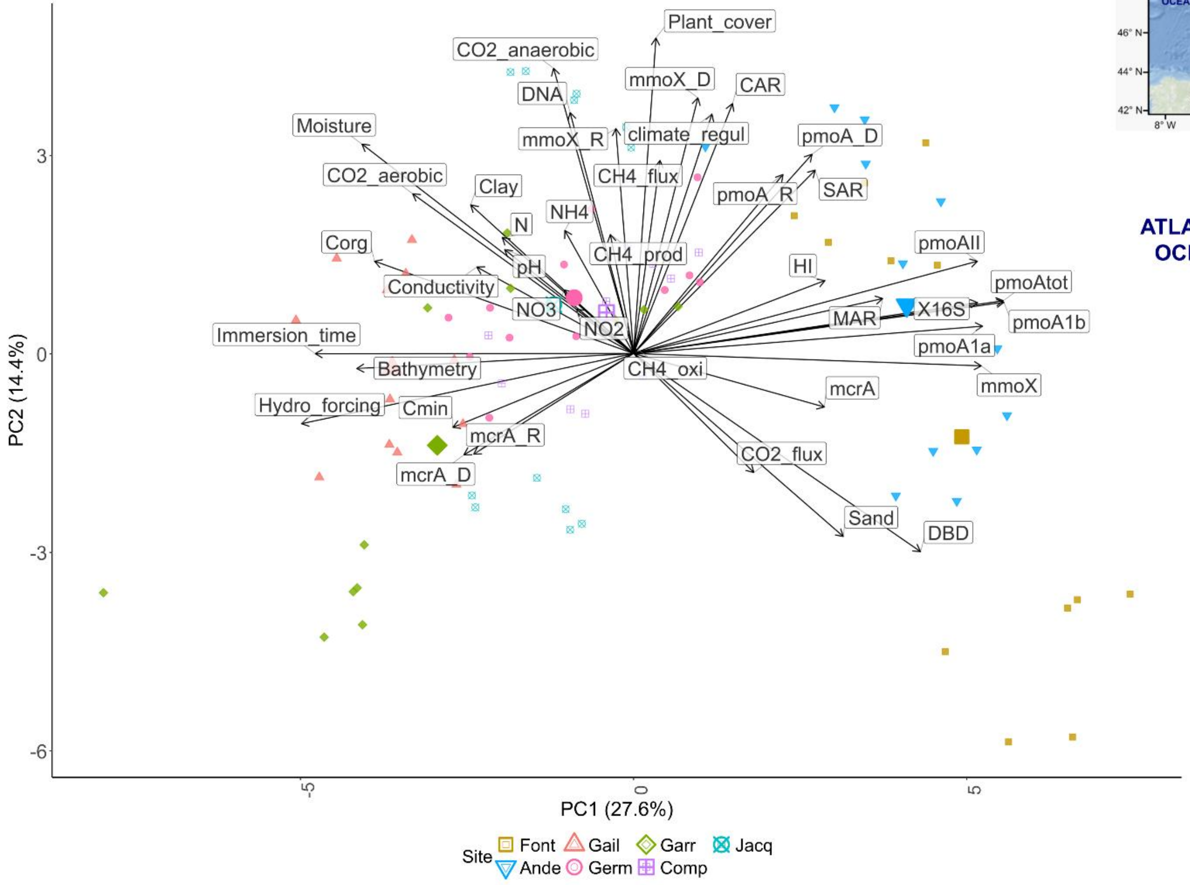

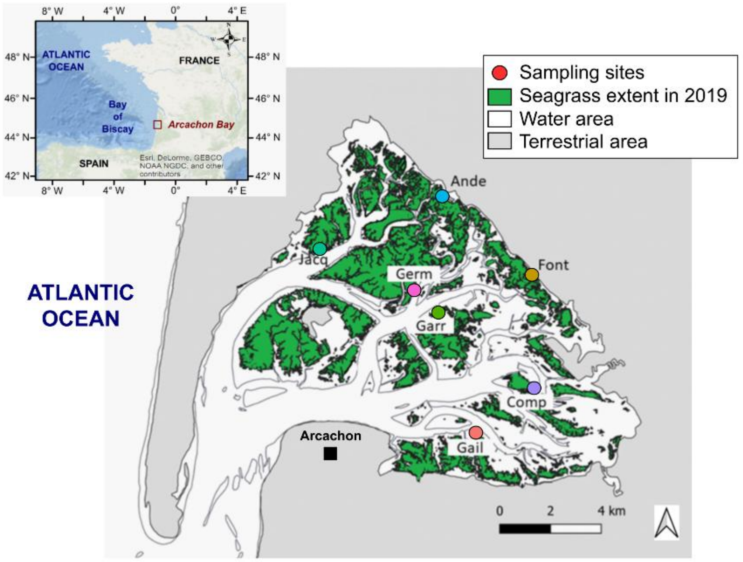

**Figure S1** Principal Component Analysis (PCA) biplot illustrating the relationships among physical, chemical, microbial, and GHG variables across seven sites (n = 83 samples), accompanied by a map of Arcachon Bay. Arrows indicate the contribution of each variable to the ordination. Both axes were rescaled (×6) to improve visualization of sites and variables. Sites are represented by different shapes and colors, with larger symbols denoting the barycenter of each site. The map uses the same color scheme for each site as the PCA plot, facilitating comparison. This supplementary figure confirms the ordination pattern shown in Figure 2 of the main text. Together, the PCA and map highlight the PC1 gradient, which reflects a large-scale ecological gradient linked to ocean connectivity and sediment stability. Sites closer to the ocean inlet, located on the left side of both figures, are characterized by longer immersion times, stronger current flows, and greater depths (higher bathymetry), such as Gail and Garr. In contrast, the sites Ande and Font, positioned on the right side of PC1, stand out as distinct and more isolated. These sites lie in the inner part of the bay, indicating a lower degree of connection to the open ocean.

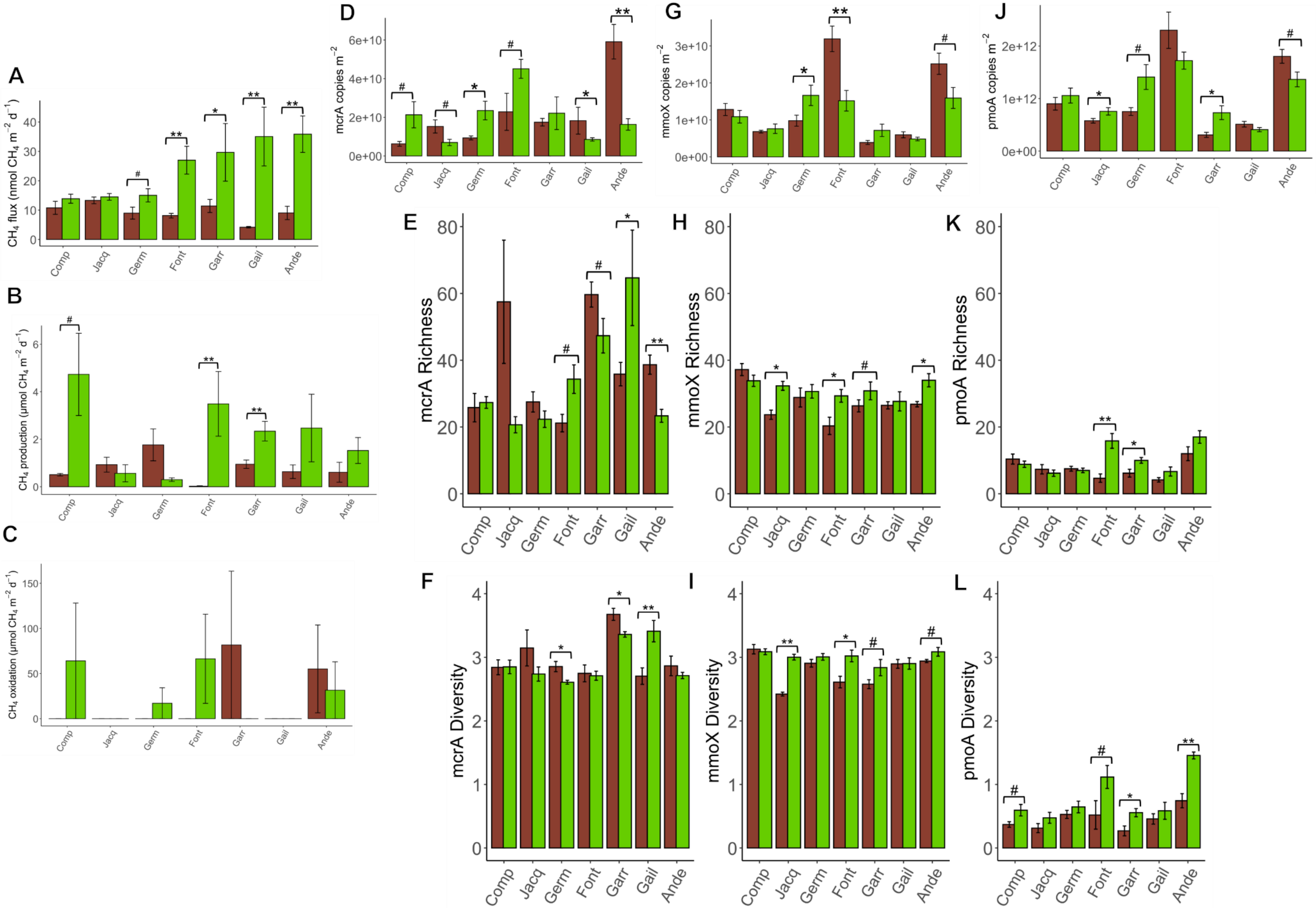
A
B
C
D
E
F
G
H
I
J
K
L
CH4 flux (nmol CH4 m-2 d-1)
CH4 production (µmol CH4 m-2 d-1)
CH4 oxidation (µmol CH4 m-2 d-1)
mcrA copies m-2
mmoX copies m-2
pmoA copies m-2
mcrA Richness
mmoX Richness
pmoA Richness
mcrA Diversity
mmoX Diversity
pmoA Diversity
Comp
Jacq
Germ
Font
Garr
Gail
Ande

**Figure S2** Comparison of $CH_4$ fluxes, $CH_4$-related processes, and microbial gene features across sites and sediment types in Arcachon Bay. Panels A-C show measured $CH_4$ flux (A), potential $CH_4$ production (B), and $CH_4$ oxidation rates (C) across sites. Panels D-F display *mcrA* gene copy numbers (D), richness (E), and diversity (F). Panels G-I show equivalent metrics for *mmoX* gene (G) copy number, (H) richness, (I) diversity. Panels J-L show *pmoA* gene copy numbers (J), richness (K), and diversity (L). Each bar represents mean ± standard error (n = 6 per subsite; n = 5 for Comp-Bare). Colors indicate sediment type: green = seagrass, brown = bare. Significant differences (Wilcoxon tests) between bare and vegetated sediments are indicated as follows: # ($p < 0.1$), * ($p < 0.05$), ** ($p < 0.01$).

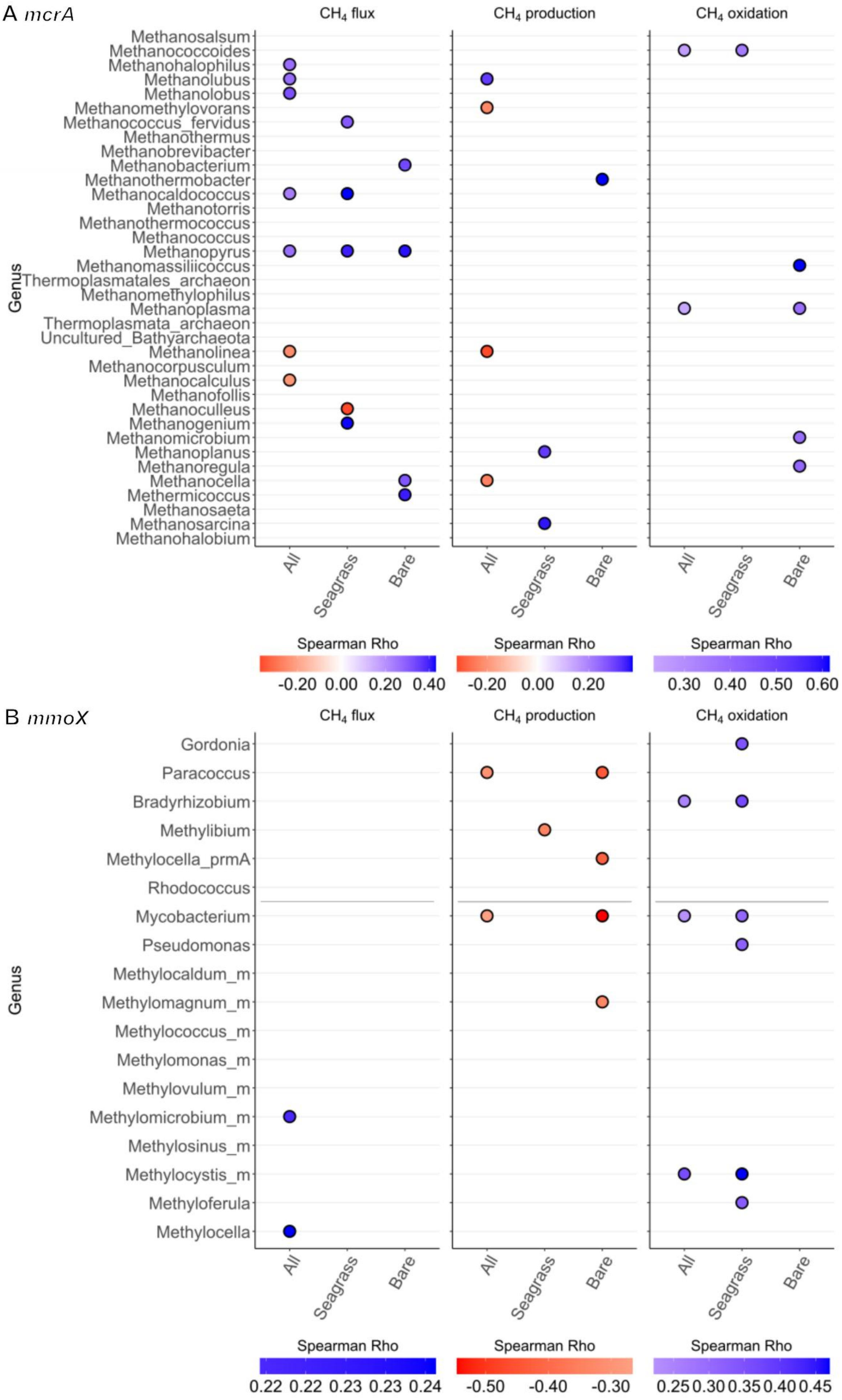
A mcrA
$CH_4$ flux
$CH_4$ production
$CH_4$ oxidation
Genus
Methanosalsum
Methanococcoides
Methanohalophilus
Methanolubus
Methanolobus
Methanomethylovorans
Methanococcus_fervidus
Methanothermus
Methanobrevibacter
Methanobacterium
Methanothermobacter
Methanocaldococcus
Methanotorris
Methanothermococcus
Methanococcus
Methanopyrus
Methanomassiliicoccus
Thermoplasmatales_archaeon
Methanomethylophilus
Methanoplasma
Thermoplasmata_archaeon
Uncultured_Bathyarchaeota
Methanolinea
Methanocorpusculum
Methanocalculus
Methanofollis
Methanoculleus
Methanogenium
Methanomicrobium
Methanoplanus
Methanoregula
Methanocella
Methermicoccus
Methanosaeta
Methanosarcina
Methanohalobium
All
Seagrass
Bare
Spearman Rho
-0.20 0.00 0.20 0.40
-0.20 0.00 0.20
0.30 0.40 0.50 0.60
B mmoX
$CH_4$ flux
$CH_4$ production
$CH_4$ oxidation
Genus
Gordonia
Paracoccus
Bradyrhizobium
Methylibium
Methylocella_prmA
Rhodococcus
Mycobacterium
Pseudomonas
Methylocaldum_m
Methylomagnum_m
Methylococcus_m
Methylomonas_m
Methylovulum_m
Methylomicrobium_m
Methylosinus_m
Methylocystis_m
Methyloferula
Methylocella
All
Seagrass
Bare
Spearman Rho
0.22 0.22 0.23 0.23 0.24
-0.50 -0.40 -0.30
0.25 0.30 0.35 0.40 0.45

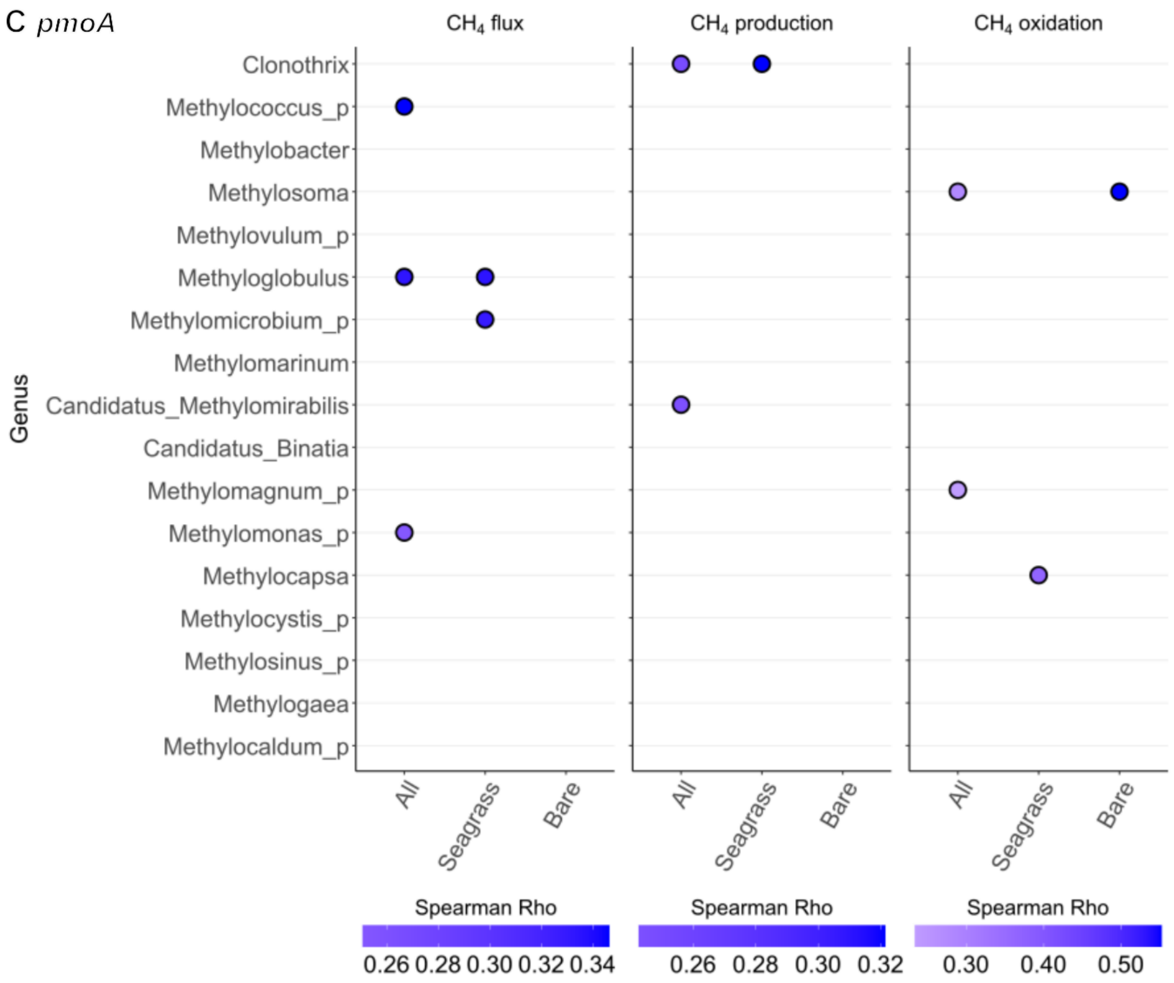

**Figure S3** Significant Spearman correlations (Rho) between genus abundances and $CH_4$-related gases. Panel A shows genera identified from the *mcrA* gene (a methanogenesis marker), panel B from *mmoX*, and panel C from *pmoA* (both methanotrophy markers). For each gene, the first column lists genera significantly correlated with $CH_4$ flux, the second column with $CH_4$ production, and the third with $CH_4$ oxidation. Colors indicate the strength and direction of the correlation (blue = positive, red = negative), n = 83.

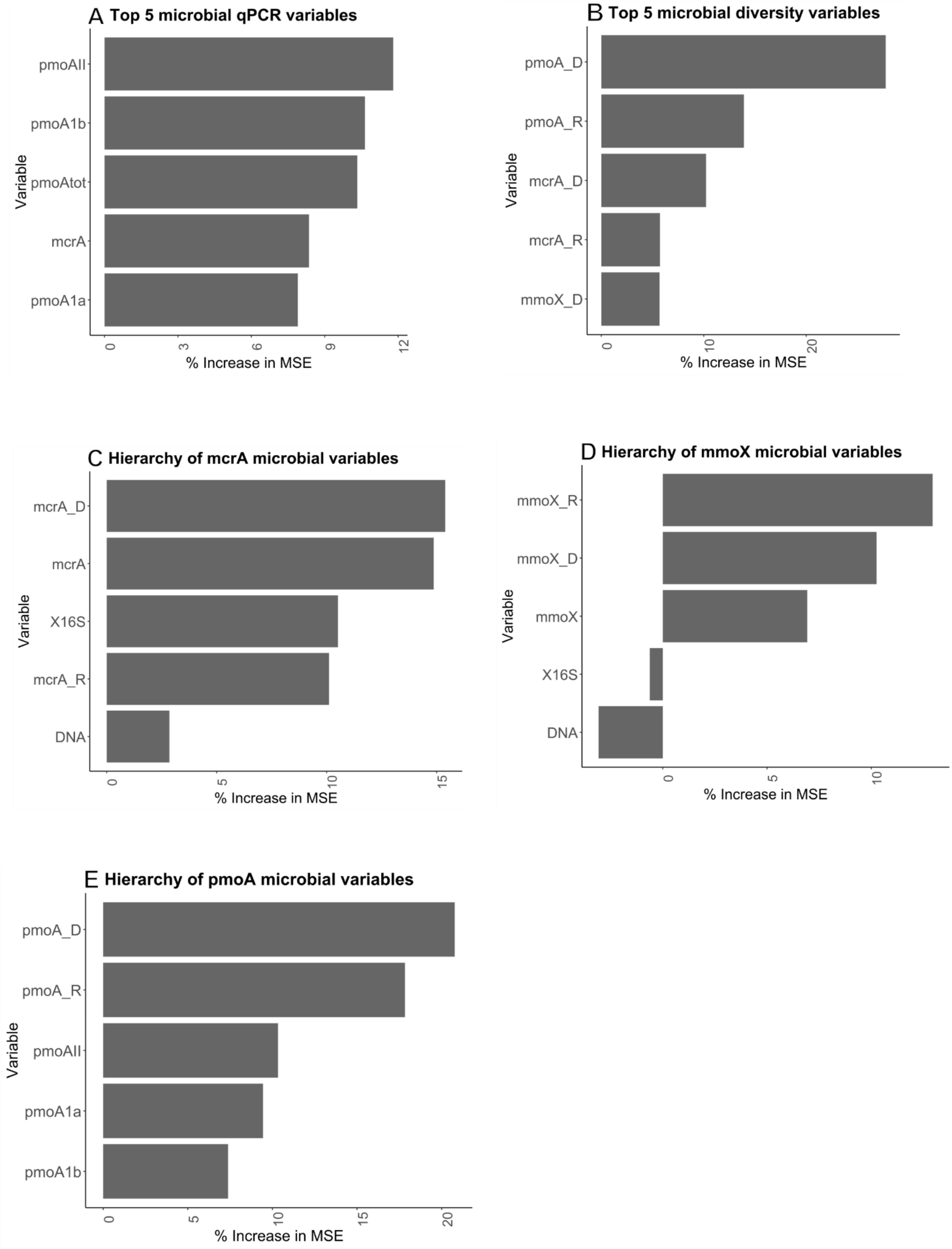
A Top 5 microbial qPCR variables
pmoAII
pmoA1b
pmoAtot
mcrA
pmoA1a
Variable
0
3
6
9
12
% Increase in MSE
B Top 5 microbial diversity variables
pmoA_D
pmoA_R
mcrA_D
mcrA_R
mmoX_D
Variable
0
10
20
% Increase in MSE
C Hierarchy of mcrA microbial variables
mcrA_D
mcrA
X16S
mcrA_R
DNA
Variable
0
5
10
15
% Increase in MSE
D Hierarchy of mmoX microbial variables
mmoX_R
mmoX_D
mmoX
X16S
DNA
Variable
0
5
10
% Increase in MSE
E Hierarchy of pmoA microbial variables
pmoA_D
pmoA_R
pmoAII
pmoA1a
pmoA1b
Variable
0
5
10
15
20
% Increase in MSE

**Figure S4** Variable importance in random forest analysis identifying key microbial predictors of methane ($CH_4$) fluxes. Importance was assessed using the mean increase in mean squared error (% Increase in MSE) after permutation of each variable, n = 83. (A-B) Top 5 predictors within each microbial category: (A) qPCR-based variables, (B) diversity indices. (C-E) Hierarchical importance of microbial predictors associated with the *mcrA*, *mmoX*, and *pmoA* genes, respectively.
